# A Statistically-Robust Model Of The Axomyelin Unit Under Normal Physiologic Conditions With Application To Disease States

**DOI:** 10.1101/2024.09.10.612208

**Authors:** Alexander Gow, Jeffrey L Dupree, Douglas L Feinstein, Anne Boullerne

**Author notes:** Correspondence: Dr Alexander Gow, Center for Molecular Medicine and Genetics, 3216 Scott Hall, 540 E Canfield Ave, Wayne State University School of Medicine, Detroit, MI, 48201.

## Abstract

Despite tremendous progress in characterizing the myriad cellular structures in the nervous system, a full appreciation of the interdependent and intricate interactions between these structures is as yet unfulfilled. Indeed, few more so than the interaction between the myelin internode and its ensheathed axon. More than a half-century after the ultrastructural characterization of this axomyelin unit, we lack a reliable understanding of the physiological properties, the significance and consequence of pathobiological processes, and the means to gauge success or failure of interventions designed to mitigate disease. Herein, we highlight shortcomings in the most common statistical procedures used to characterize the axomyelin unit, with particular emphasis on the underlying principles of simple linear regression. These shortcomings lead to insensitive detection and/or ambiguous interpretation of normal physiology, disease mechanisms and remedial methodologies. To address these problems, we syndicate insights from early seminal myelin studies and use a statistical model of the axomyelin unit that is established in the accompanying article. Herein, we develop and demonstrate a statistically-robust analysis pipeline with which to examine and interpret axomyelin physiology and pathobiology in two disease states, experimental autoimmune encephalomyelitis and the *rumpshaker* mouse model of leukodystrophy. On a cautionary note, our pipeline is a relatively simple and streamlined approach that is not necessarily a panacea for all *g* ratio analyses. Rather, it approximates a minimum effort needed to elucidate departures from normal physiology and to determine if more comprehensive studies may lead to deeper insights.

## INTRODUCTION

In the decades following the pioneering work on PNS myelinated fiber morphology by Donaldson and Hoke (1905), relationships between Schwann cell-derived compact myelin and the ensheathed axons were studied extensively. Conduction velocity (CV), myelin thickness, axon diameter and internodal length were found to exhibit linear (or close-to-linear) correlations across a broad range of vertebrate species (reviewed by Rushton, 1951). Common practice was to represent these associations as relations of internodal length or fiber diameter, particularly for the *g* ratio (defined as the ratio of axon diameter to fiber diameter).

Such practices led to a rigorous theoretical model (Rushton, 1951) that underpins much of our understanding about myelin function, its structural and physiological properties and computational modeling. Arguably, the most important morphological and functional properties of a robust axomyelin unit model in large diameter (> 3μm) PNS fibers were: (1) strong dependences of CV on fiber diameter and internodal length, (2) weak dependence of CV on the *g* ratio for a given fiber diameter (in the observed range, 0.5 < *g* ratio < 0.8), and (3) independence of the *g* ratio on fiber diameter.

For small PNS and CNS myelinated axons (< 3μm), there was considerable doubt for decades that *g* ratio was independent of fiber diameter. However, this notion was finally dispelled when the flaws in measurement methods of came to light. Thus, limit-of-resolution (Rayleigh diffraction limit) errors in brightfield micrographs overestimated the myelin thickness (thereby decreasing *g* ratio), with the error inversely proportional to fiber diameter (Schnepp and Schnepp, 1971). Subsequently, the anticipated independence of *g* ratio on fiber diameter was established from electron micrographs for small fibers in multiple species and white matter tracts (reviewed in Waxman and Swadlow, 1977).

Easily-measured properties that established morphologic (*g* ratio) and functional (CV) relationships for single large fibers in PNS nerves and spinal roots (reviewed in Rushton, 1951) were difficult to replicate in the CNS, except for a few studies in large diameter spinal cord myelinated fibers and computer simulations based largely on the properties of peripheral fibers (Blight and Someya, 1985). Nevertheless, Waxman & Bennett (1972) bridged this gap by refining the theoretical predictions of Rushton (1951) and computed CVs as a relation of myelinated fiber diameter for the smallest CNS fibers. These estimates, which indicate CV is slower for myelinated versus unmyelinated axons of ≈0.2μm or less in diameter, accounts for the observed minimum caliber of myelinated axons in the CNS. Further, they are consistent with CV measurements derived from optic nerve compound action potentials (CAPs) and several spinal tracts, as well as more recent in silico models (Devaux and Gow, 2008; Gow and Devaux, 2008).

The importance of CV, myelin thickness, internodal length and fiber diameter relationships notwithstanding, Gasser and Grundfest (1939) used the CAP reconstruction method (Gasser and Erlanger, 1927) to advocate for shifting focus toward correlations of CV, *g* ratio and myelin thickness with axon diameter under disease states (e.g. multiple sclerosis, neuropathies and the leukodystrophies), and away from the uncorrelated *g* ratio versus fiber diameter relation. This emphasis persists in contemporary studies, so much so that increases in *g* ratio have become synonymous with decreases in CV, which is inconsistent with theoretical predictions (Rushton, 1951). Moreover, there is heightened risk for this perspective to be overstated or misinterpreted, not least because of the incorrect statistical methods, such as simple linear regression, that are typically used for data analysis.

In this light, we revisit the original notion of the *g* ratio as a relation of fiber diameter (Donaldson and Hoke, 1905), where a formal adoption of the uncorrelated properties of these two parameters permits robust representation of the axomyelin unit. Sagacious use of the statistical model thereby afforded, yields outcomes that are relatively free of technical artifacts (see accompanying article). Herein, we demonstrate implementation of an analysis pipeline for the axomyelin unit under normal physiological conditions, with implications for two disease states from previously published reports (Dupree et al., 2015; Southwood et al., 2017). Automation of this pipeline is currently underway in the Graphpad Prism and Excel software platforms.

## METHODS

### Experimental data and assurances of humane care of mice

All experimental data from mice that are presented in the current project were obtained from previously published studies (Dupree et al., 2015; Southwood et al., 2017). The authors stated in those publications, and we concur, that all animal procedures were humane and approved by the Institutional Animal Care and Use Committees from the respective institutions.

### *g* ratio measurement in the experimental autoimmune encephalomyelitis (EAE) mouse model of multiple sclerosis

Raw data for the current study were obtained from optic nerves of adult Sham-treated mice analyzed previously by Dupree and colleagues (2015). Briefly, experimental autoimmune encephalomyelitis (EAE) was induced in mice using MOG_35-55_ peptide in saline emulsified in complete Freund’s adjuvant, including *M. tuberculosis* and pertussis toxin. A Sham-treatment cohort received the same EAE-inducing cocktail and regimen as the EAE cohort except for the MOG_35-55_ peptide. Seven weeks later, corresponding to 40 days after peak clinical disease scores, mice were perfused with 4% paraformaldehyde / 5% glutaraldehyde in 0.1M Millonig’s buffer and processed into PolyBed 812 resin for electron microscopy. Ultrathin plastic sections of transverse optic nerve were mounted onto standard 200 mesh uncoated copper grids, and all correctly-oriented (normal to the plane of section) myelinated fibers were surveyed in 10 randomly selected fields (magnification: 10,000x; camera pixel resolution = 2.27nm) per mouse. Axon and fiber diameters were measured digitally (the average between minimum and maximum diameter measurements) for plots and statistics. Unmyelinated axons, and fibers cut obliquely to the long axis were not included in the current study.

### *g* ratio measurement in the *rumpshaker* (*rsh)* mouse model of Pelizaeus Merzbacher disease

The raw data for the current study were obtained from 10 week old mouse optic nerves analyzed previously by Southwood and colleagues (2017). The circumference and radial width of compact myelin (at a location where the major and minor dense lines were visible) for each myelinated axon were captured on photographic negatives at magnifications of 10,000x (large fibers) or 15,000x (small fibers). The negatives were scanned at 1200 d.p.i. (final pixel resolution = 2nm) for measurements using a 9” x 12” digital tablet (model XD-0912-U, Wacom Co. Ltd.). Equivalent axon diameters were computed from the circumference, and divided by fiber diameter (sum of the axon diameter and twice the myelin width) to obtain the *g* ratio. Width of the periaxonal space was assumed to partition equally between the axon diameter and myelin width measurements (i.e. within the experimental errors) and was not explicitly added to compute fiber diameter. Unlike the EAE study, myelinated fibers measured in the *rsh* study were not uniformly derived from random fields. Rather, a preponderance of high quality large diameter fibers were selected to ensure a broad range of fiber diameters for analysis.

### Independent versus pseudo-replicate *g* ratio measurements and the grand average

To determine *g* ratios in myelinated fiber tracts, 100-150 individual axons are measured from each mouse and compiled in a scatterplot. Historically, many groups have regarded measurements from individual animals as independent *g* ratio measurements; however, they are actually pseudo-replicate data (measurements from a single animal under identical experimental conditions) because they are subject to systematic experimental errors. By definition, independent data are subject only to random experimental errors. As examples of systematic errors, if the perfusion fixative were incorrectly prepared or if the perfusion was of low quality, the morphology of all axons in the mouse would be distorted to a similar extent. A simple approach to avoid problems with replicates is to compute the grand average *g* ratio per mouse (a single estimate per mouse). From a *g* ratio versus fiber diameter scatterplot for each cohort, the median *g* ratio per mouse is computed. These medians are then averaged to generate the grand average (mean ± standard deviation (S.D.) of the cohort for statistical testing using t-tests or mixed-effects analysis (similar to repeated measures ANOVA).

### *g* ratio analysis kernel and pipeline

The analysis pipeline comprises a large series of steps and decision points, but the basic kernel is summarized in Table 1. Beginning with the raw data measured from electron micrographs at a pixel resolution no greater than 3nm, measurements of axons (min/max diameters or circumferences) and fibers (min/max diameters, circumferences or 2 x radial myelin thickness = myelin diameter) are transformed to obtain the equivalent axon and fiber diameters. Axon diameter as a relation of fiber diameter is plotted and fit using linear regression to determine if the relation is directly proportional (i.e. linear and passing through the Origin within the margin of error). Visual inspection of the plot is an important quality control because linearity and strong correlation (i.e. R^2^ ≅ 1) in the cohort indicates there are no major artifacts with data collection (e.g. limit of resolution artifacts).

**Table 1.**
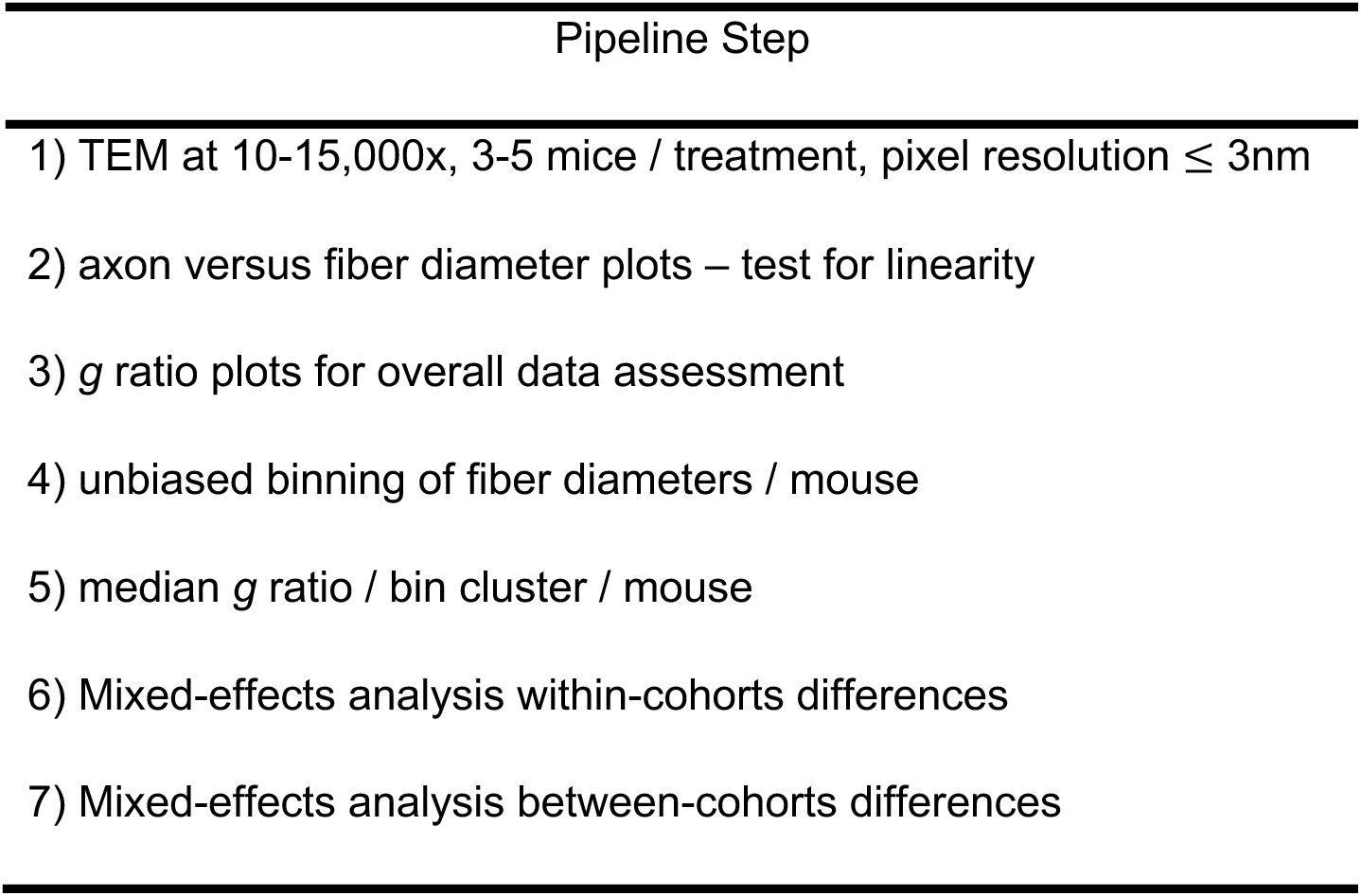
the analysis kernel for the *g* ratio pipeline.

Next, *g* ratio histograms are generated for each mouse from an ordered list of fiber diameters apportioned into arbitrarily-defined ranges (fiber diameter bins or clusters). Fiber diameter ranges are chosen from one mouse so that each cluster contains approximately 20 *g* ratios. Accordingly, the number of clusters might vary according to the number of *g* ratios per mouse (e.g. 120 *g* ratios equates to 6 clusters). The same fiber diameter ranges are used for the remaining mice in the experiment without concern for the exact number of *g* ratios in each cluster. The purpose of establishing consistent fiber diameter ranges is to ensure that direct comparison is possible for statistical analysis.

Visual inspection of the resulting histograms from the cohorts is an important quality check to ensure *g* ratios are unimodal with normal or slightly skewed distributions in the majority of the clusters. Medians [±95% confidence intervals (CI)] are used to summarize the clusters, which minimizes the impact of small deviations from normally-distributed data. Confidence intervals provide the advantage of a visual check on data quality; the grand average should be within the 95% CI of most clusters if independence assumption between *g* ratio and fiber diameter is met (for subsequent steps in the pipeline, computing means ±S.D. is suitable).

At this point of the pipeline, it is tempting to automate the visual inspection of clustered data using normality tests such as D’Agostino-Pearson omnibus K2, Shapiro-Wilk, Anderson-Darling or Q-Q plots. However, such approaches are often unreliable and insensitive for small sample sizes below 30-50. Moreover, a major assumption underlying these tests is that all data points are independent measurements. Their use is not statistically justified with the pseudo-replicate data generated for *g* ratio plots.

### Hypomyelination-hypermyelination quartiles for the *g* ratio scatterplot

Horizontal shaded quartile zones in scatterplots are computed from the grand averages of the control cohort, using the following relation:

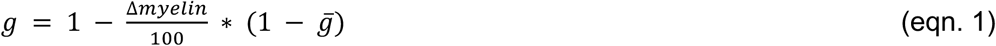

where *g̅* is the grand average for the control cohort, Δmyelin is a prescribed percentage of myelin in the mice that may be lost or gained (±25%, ±50%, ±75%), and *g* delineates the ordinate values corresponding to the shaded zones of myelin loss or gain.

### Statistics methods to ensure unbiased analyses

Microsoft excel (ver16.66.1) was used to collate raw data measurements from TEM micrographs (Dupree et al., 2015; Southwood et al., 2017) and compute axon and fiber diameters, and *g* ratios. These data were copied into Graphpad Prism v10.2.1 (GraphPad Software, LLC, San Diego, CA) for all subsequent processing steps and statistical analyses. Results of statistical tests are reported in the figure legends. Adjusted P values were computed for all multiple comparison tests. Mixed-effects analysis with Geisser-Greenhouse epsilon correction and Tukey’s multiple comparison’s post hoc test was used to account for within-subjects pseudo-replicate correlations (i.e. systematic errors). Mixed-effects analysis with Geisser-Greenhouse epsilon correction and Sidak’s multiple comparisons post hoc test was used for between-treatments testing. Box-Cox and Fisher’s Z transformations were used to demonstrate the linearity of axon-fiber diameter relations in all cohorts (see accompanying article, Figs S2-5).

### Deming regression versus simple linear regression

Virtually all myelin internode studies over the last few decades use simple linear regression (least squares regression) to analyze myelin and *g* ratio plots. An implicit assumption with this approach is that experimental errors are only associated with the dependent variable, while errors in the independent variable are assumed to be negligible/absent. This assumption is problematic because experimental errors are present in both axon and fiber diameter measurements, so the appropriate analysis is Deming linear regression (i.e. orthogonal linear regression), which we use to analyze axon versus fiber diameter plots.

Deming linear regression has limitations when used with correlation plots (i.e. if experimental errors are present in the correlated variables), and cannot be used to perform inferential statistics between mice because the magnitude of standard errors and confidence intervals are too small, thereby increasing type I errors. The regression approach can be used with caution for exploratory analyses such as descriptive statistics, visualization, R^2^ goodness of fit estimates of correlation between experimental measurements and informal hypothesis generation.

## RESULTS

Use of *g* ratios to assess compact myelin from PNS and CNS diseases in multiple species including humans is common, widespread and arguably, the gold standard for analysis. A number of laboratories report *g* ratios as a single value per animal (i.e. the grand average), but the typical method is the *g* ratio scatterplot. Almost universally, these plots comprise 100-200 measurements per animal and 3-4 animals per treatment group. The *g* ratios are plotted as a relation of axon diameter and analyzed using simple linear regression. However, this approach has major technical drawbacks associated with inappropriate statistics, and conceptual deficits because of the lack of an explicit model of the axomyelin unit with which to constrain the interpretation of the analyses.

To remedy our broad unease about such free-wheeling analyses, the accompanying article establishes an axomyelin unit model that is used in a data processing pipeline described herein (Fig. 1). The pipeline kernel is summarized in Table 1, and two case studies using the pipeline are detailed below. The key advance over previous analyses is an explicit set of model assumptions that serve as a framework for constraining the interpretation of *g* ratio data. Indeed, these assumptions provide clear outcomes and testable hypotheses. In furtherance of our objectives, we draw on published *g* ratio data from two studies to illustrate our approach.

**Figure 1.**
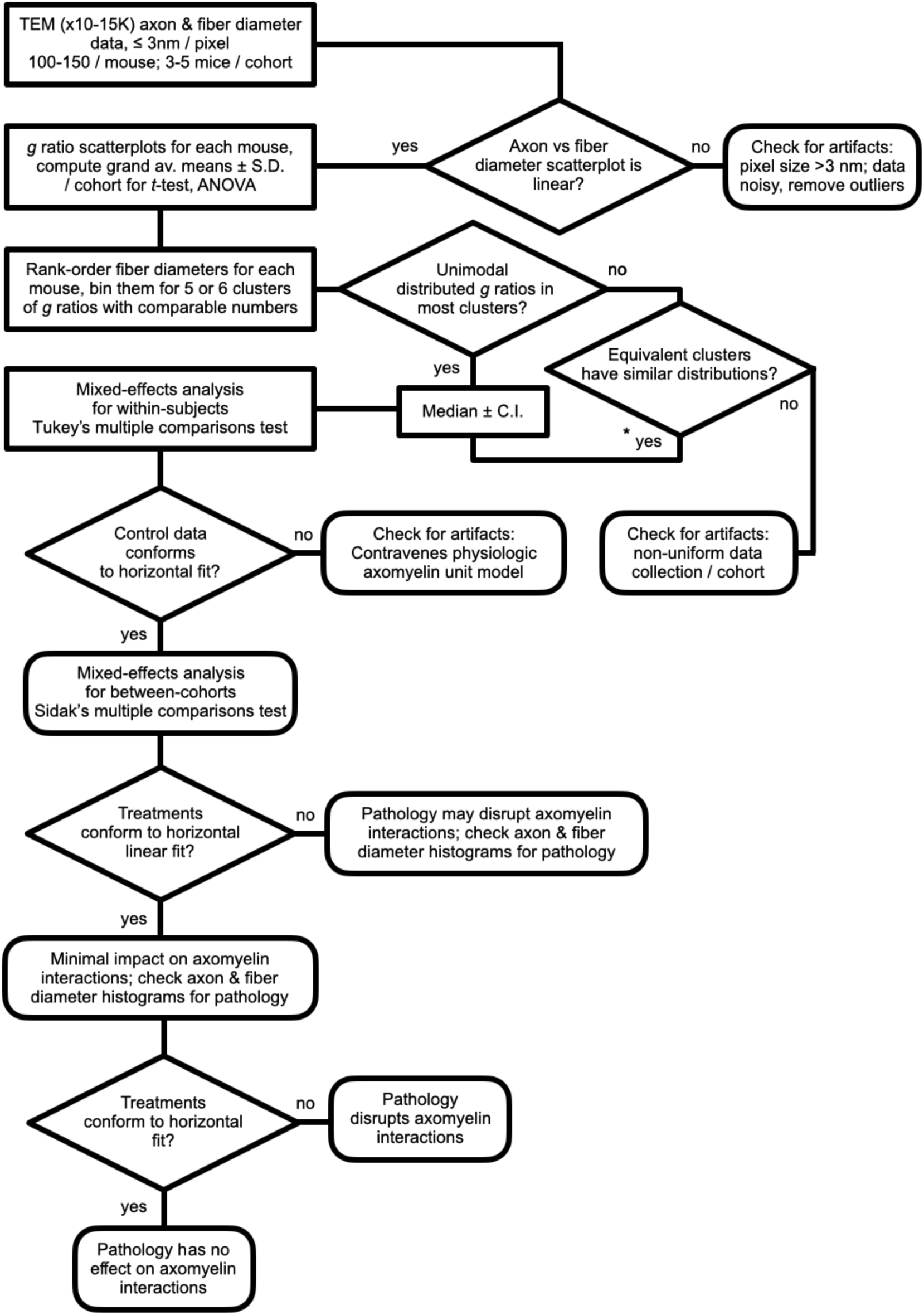
pipeline and decision tree to analyze *g* ratios from T.E.M. data: a robust model for the axomyelin unit. Beginning with transmission electron micrographs (bright field micrographs are not appropriate for measurements of myelinated fibers < 3μm), measure axon diameters (or circumference) and fiber diameters (or circumference) or myelin thickness for 100 – 150 fibers per animal. Compute equivalent diameters from circumferences as necessary. Paste the *g* ratios and fiber diameters into a Graphpad or excel template to complete the analysis. **\*** Even if equivalent *g* ratio clusters (i.e. the same fiber diameter bin) are similar for each mouse in a cohort, summarizing bimodal and other non-unimodal distributions may yield questionable results. Such data arising in control cohorts may signify variable data collection or analysis methods and the error source should be identified and eliminated. Non-unimodal distributions that emerge from test cohorts also may reflect variable data collection methodology (e.g. by including unmyelinated fibers, where *g* ratio = 1), but may indicate multiple pathological processes.

### Analysis of the axomyelin unit under physiological conditions

To develop a pipeline for analysis of the axomyelin unit, we first used data from optic nerves of Sham-treated adult mice published by Dupree and colleagues (2015). These data also are used in the accompanying article, reproduced in Fig. 2A for the reader’s convenience, to establish an axomyelin unit model and identify several common artifacts in *g* ratio plots. The grand average *g* ratio plot is also reproduced, in Fig. 2B. While statistically-rigorous because it eliminates systematic errors (i.e. pseudo-replicate), complete reduction of data to a single mean ±S.D. discards most of the morphological details gleaned from electron micrographs, and can reduce the sensitivity of statistical tests to detect subtle pathophysiological changes in disease states.

**Figure 2.**
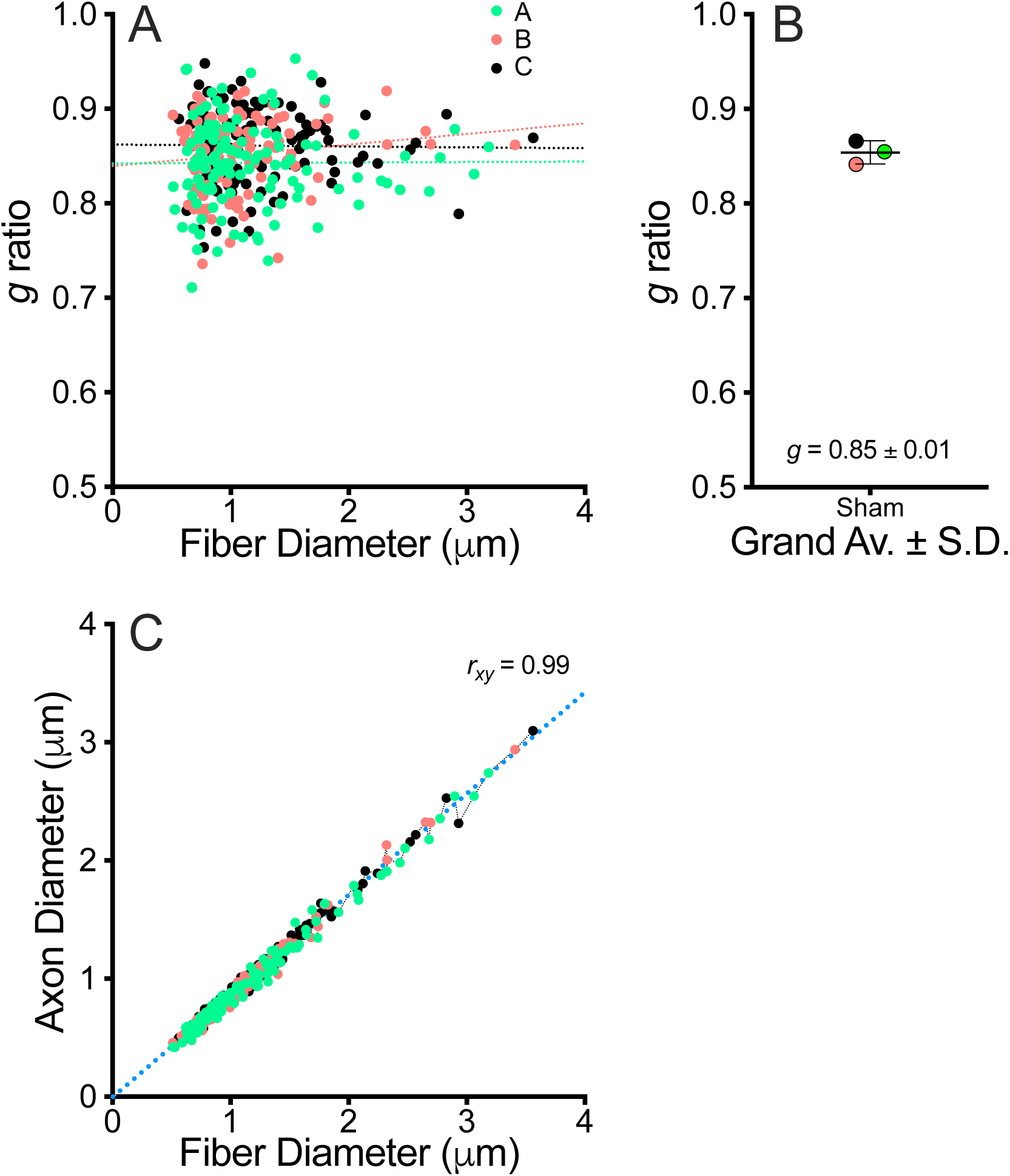
optic nerve *g* ratio versus fiber diameter scatterplot from Sham mice. **A.** *g* ratio scatterplot from transmission electron micrographs for three Sham mice, A-C. Simple linear regression fits are similar for each mouse, but regression slopes and *y*-intercepts are unstable. Most of the data are clustered between fiber diameters of .5 – 2μm and *g* ratios between .6 – .9. **B.** The median *g* ratio for each mouse and the grand average for the cohort, .85 ± .01 (mean ±S.D.) are computed from the scatterplots in (A). **C.** Correlation plot between axon and fiber diameters indicates a very strong linear relation (*r_xy_* = .99). A Deming linear regression fit (blue dotted line) passes through the Origin (*y*-intercept = −.009, 95% CI [−.024, .005]), indicating direct proportionality between the two variables.

From a statistical modeling standpoint, the commonly published method of simple linear regression fits to *g* ratio plots is incongruous for several reasons (Table 2). First, the approach discounts a fundamental assumption of simple linear regression, that only random errors are permitted sources of variance in experimental data. Measurements for *g* ratio analysis are pseudo-replicates (not independent data points). Second, a regression fit assumes data are uniformly distributed across the independent and dependent variable domains. But, much of the data in Fig. 2A are clustered (between fiber diameters of 0.8-2μm and *g* ratios of 0.6-0.9, Fig. S1A-C), with relatively few measurements outside this central region. Such regions of sparce data may include influential data points that disproportionately distort regression fits.

**Table 2.** false assumptions associated with using simple linear regression in *g* ratio studies.

| False Assumption |
| --- |
| 1) simple linear regression is appropriate for *g* ratio studies |
| 2) pseudo-replicate data are compatible with simple linear regression in statistics tests |
| 3) *g* ratios are directly correlated with axon diameters |
| 4) every axon-fiber diameter measurement in each animal is an independent observation |
| 5) measurements need not be evenly spread along the x-axis |
| 6) *g* ratios need not be normally distributed across the regression curve |
| 7) regression fits for *g* ratio plots have non-zero slope |

A third incongruity is that the requirement of a uniform and random distribution of the dependent variable along the regression line (called homoscedasticity) is not met. Finally, the regression fits in Fig. 2A have non-zero slopes, implying a correlation between *g* ratio and fiber diameter. The accompanying article demonstrates that no such correlation exists in this dataset, which is consistent with the early literature (Donaldson and Hoke, 1905; Rushton, 1951; Schnepp and Schnepp, 1971; Price and Sprich, 1975; Waxman, 1980). Thus, current practices contravene decades of rigorous experimental studies in multiple vertebrate species.

These incongruities notwithstanding, plots of raw data from electron micrographs (high Pearson’s correlation coefficient, Fig. 2C) are themselves the strongest indication that *g* ratio plots can be misleading. Indeed, the Box-Cox analysis in Fig. S2 (reproduced from the accompanying article for convenience) demonstrates the axon-fiber diameter relation is linear using statistics tests and accords with expected internodal biology. Moreover, the observation that the Deming regression fit to the axon-fiber diameter plot (Fig. 4C) passes through the Origin (*y*-intercept = −0.004, 95% CI: [−0.017, 0.008]) demonstrates the direct proportionality of the relation and prescribes that *g* ratio plots conform to a horizontal regression fit (approximating the grand average *g* ratio), as predicted by the axomyelin unit model.

### Balancing statistical rigor and information content

In view of the artifacts associated with typical *g* ratio plots (see accompanying article), the pursuit of alternate strategies to summarize axomyelin unit properties, satisfy major statistical assumptions and preserve information content is of primary importance. Our analysis pipeline (Fig. 1) implements a series of processing steps and statistical tests to accomplish this task. Thus, fiber diameters from each mouse are rank-ordered, and the *g* ratios are clustered (binned) with similar numbers of *g* ratios per cluster. Binning the *g* ratios minimizes systematic effects of pseudo-replicate data and reduces the initial number of data points for each mouse, which improves the validity of subsequent statistical tests. Each cluster is labeled according to the maximum fiber diameter boundary, and constituent *g* ratios are represented as frequency histograms (Fig. 3A).

**Figure 3.**
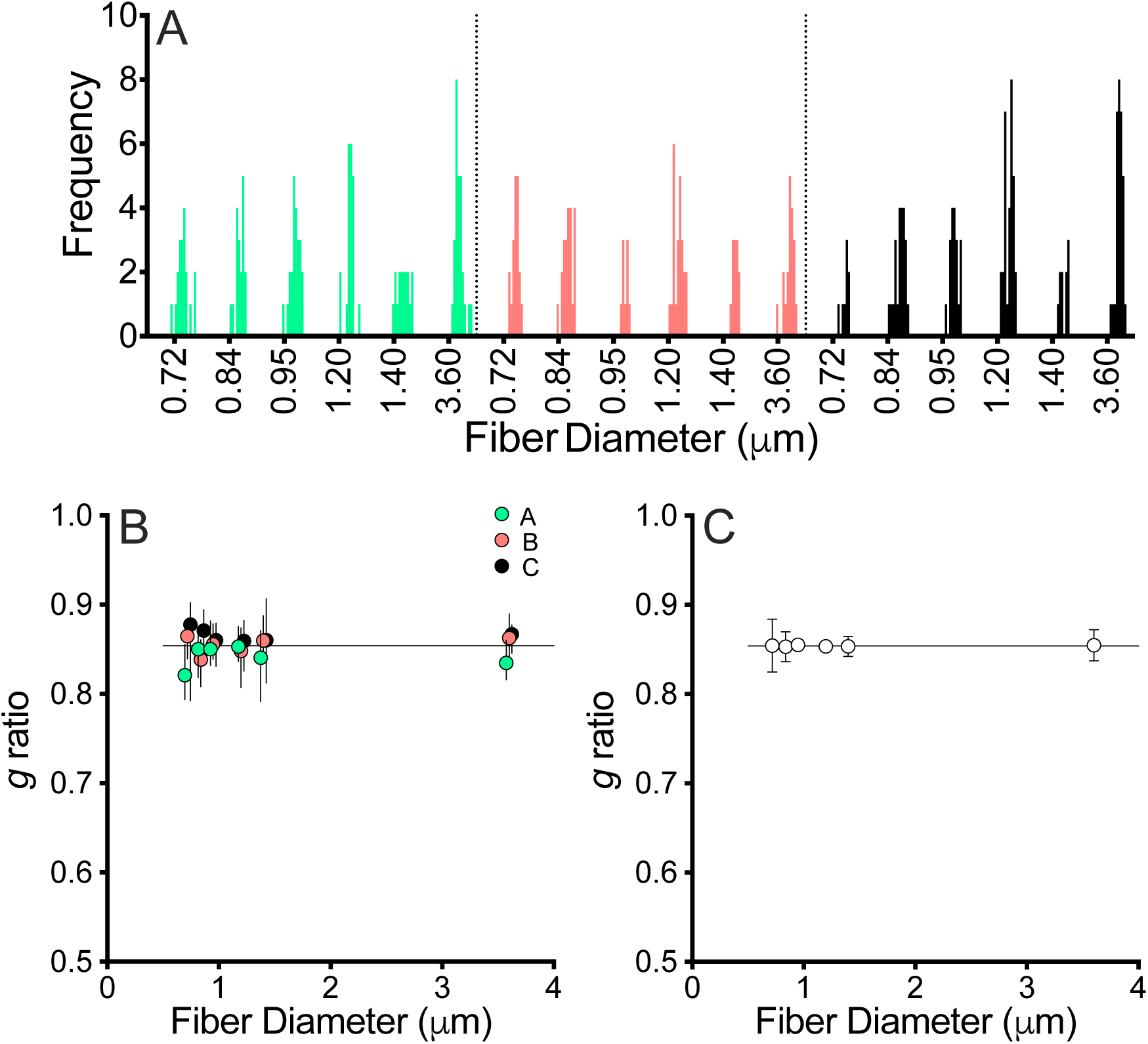
*g* ratio scatterplot quality control and data reduction for Sham treated mice. Scatterplot data from mice A-C (from Fig. 2A) sorted by fiber diameter (ascending order) and binned into six clusters with similar numbers of *g* ratios per cluster (the same cluster boundaries are used for each mouse). **A.** Histograms show *g* ratios in most clusters are unimodal with normal or skewed distributions and can be summarized as medians ±95% CI for each mouse. **B.** The summary data plotted as a relation of the maximum fiber diameter boundary in each cluster for comparison with the horizontal line (grand average for the cohort, Fig. 2B). The data for mice A and C are shifted left and right on the *x*-axis, respectively, to show all the 95% CIs. **C.** Corresponding cluster means from each mouse are averaged (mean ±S.D.). Mixed-effects analysis with Geisser-Greenhouse correction indicates there are no significant differences between clusters (F_(1.48,2.96)_ = .014, ε = .30, P = .97), indicating *g* ratio is uncorrelated with fiber diameter. This result is consistent with the direct proportionality of the axon-fiber diameter relation (Fig. 2C).

Three features are salient. First, six fiber diameter clusters containing the *g* ratios (this number can be higher or lower) are defined for each mouse. Second, *g* ratios within the clusters are unimodal and have normal or slightly skewed distributions, indicating that medians are the appropriate summary statistic. These cluster medians are plotted ±95% CI in Fig. 3B. A horizontal line equal to the cohort grand average = 0.85 (Fig. 2B) passes through the 95% CIs of the *g* ratio medians, which validates this method of summarizing the *g* ratio data. In Fig. 3C, averaging the medians of corresponding clusters (mean ±S.D.) for mixed-effects analysis indicates there are no differences between cluster means. Thus, *g* ratio clusters accord with the grand average *g* ratio, and we conclude that *g* ratio is uncorrelated with fiber diameter per the axomyelin unit model.

### Autoimmune demyelination has marginal impact on *g* ratios from optic nerve

Dupree and colleagues (2015) measured optic nerve *g* ratios in mice 40 days after peak clinical disease scores. These data are plotted as a relation of fiber diameter in Fig. 4A. The overall data structure is comparable to the Sham cohort (Fig. 2A) including instability of the regression slopes between mice. Nevertheless, the correlation between axon and fiber diameters in individual mice is strongly linear (Figs 4B, S3), similar to the controls (Figs 2C, S2), and a Deming regression fit passes through the Origin (*y*-intercept = −0.004, 95% CI: [−0.017, 0.008]) indicating a direct proportionality relation. Comparison of the grand averages (Fig. 4C) for the Sham and EAE groups shows there is no statistical difference, which was the conclusion reached in the original study of these mice (Dupree et al., 2015).

**Figure 4.**
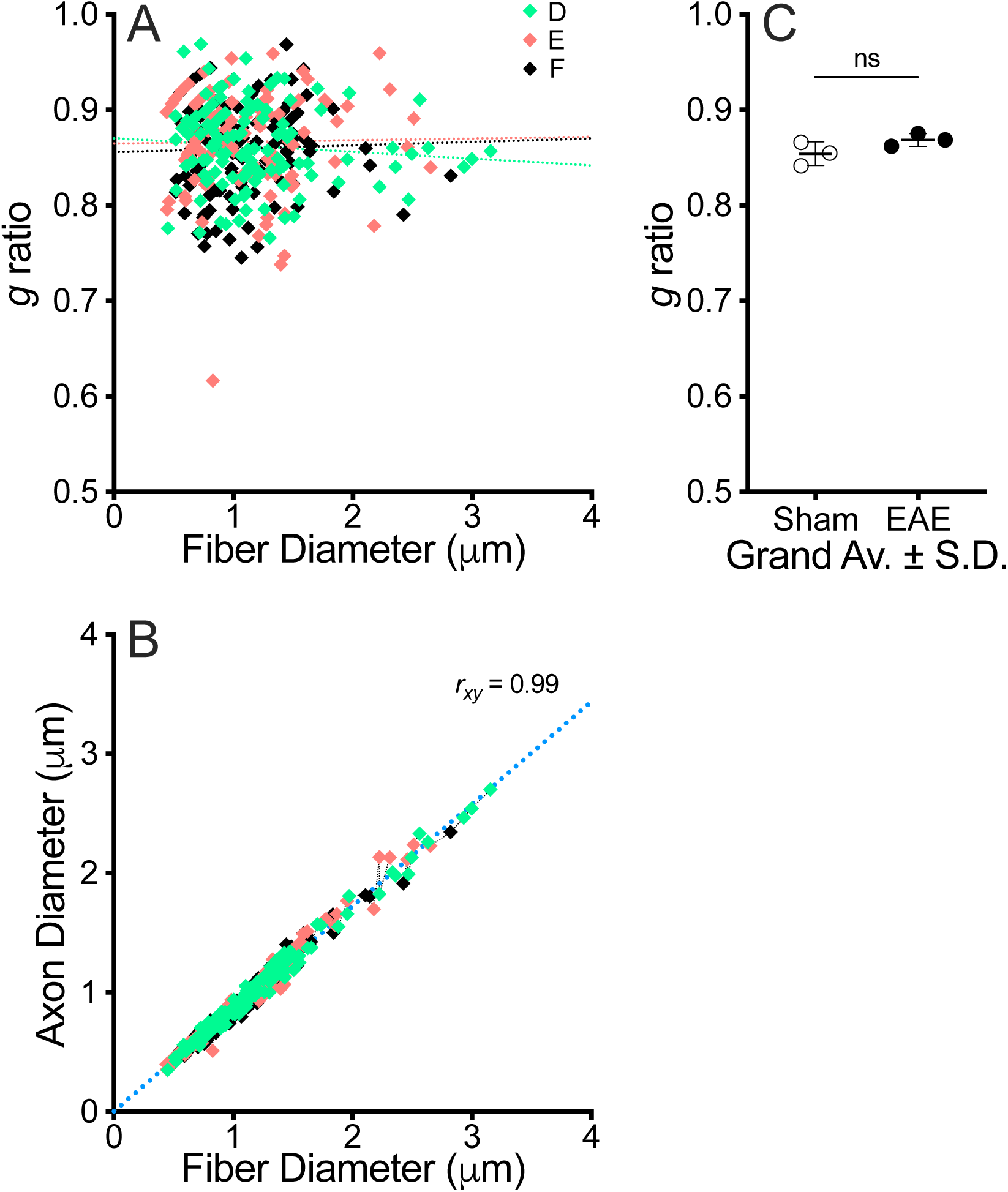
optic nerve *g* ratio versus fiber diameter scatterplot from EAE mice. **A.** *g* ratio scatterplot from transmission electron micrographs for three EAE mice, D-F. The slope components of simple linear regression fits are unstable. Most of the data are clustered between fiber diameters of .5 – 2μm and *g* ratios between .75 – .95. **B.** Correlation plot between axon and fiber diameters indicates a very strong linear relation (*r_xy_* = .99). A Deming linear regression fit (blue dotted line) passes through the Origin (*y*-intercept = −.002, 95% CI [−.018, .014]), indicating direct proportionality between these two variables. **C.** The median *g* ratio for each mouse and the grand average for the cohort, .87 ± .01 (mean ±S.D.) are computed from the scatterplot in (A). The grand average from the Sham treated mice (Fig. 2B) is included for comparison. An unpaired t-test indicates the *g* ratios are indistinguishable (P = .14, *n* = 3 per cohort).

Clustering *g* ratios from the EAE mice (equal to cluster boundaries in Fig. 3A) shows the histograms are unimodal with normal or skewed distributions (Fig. 5A). Summarizing these data as medians ±95% CI (Fig. 5B) indicates the clusters are consistent with the grand average for the EAE cohort. Means ±S.D. are shown in Fig. 5C, and a mixed-effects analysis indicates *g* ratio is not correlated to fiber diameter (horizontal regression fit). Thus, any potential myelin pathology would be relatively consistent across the vast majority of myelin internodes. Combining the Sham (Fig. 3C) and EAE *g* ratio cluster mean ±S.D. summaries in Fig. 5C shows the *g* ratio clusters from these treatment groups are similar and a mixed-effects analysis indicates no differences between the treatment groups. Thus, our analysis pipeline confirms the original results (Dupree et al., 2015) but yields deeper insight into the effects of EAE in optic nerve.

**Figure 5.**
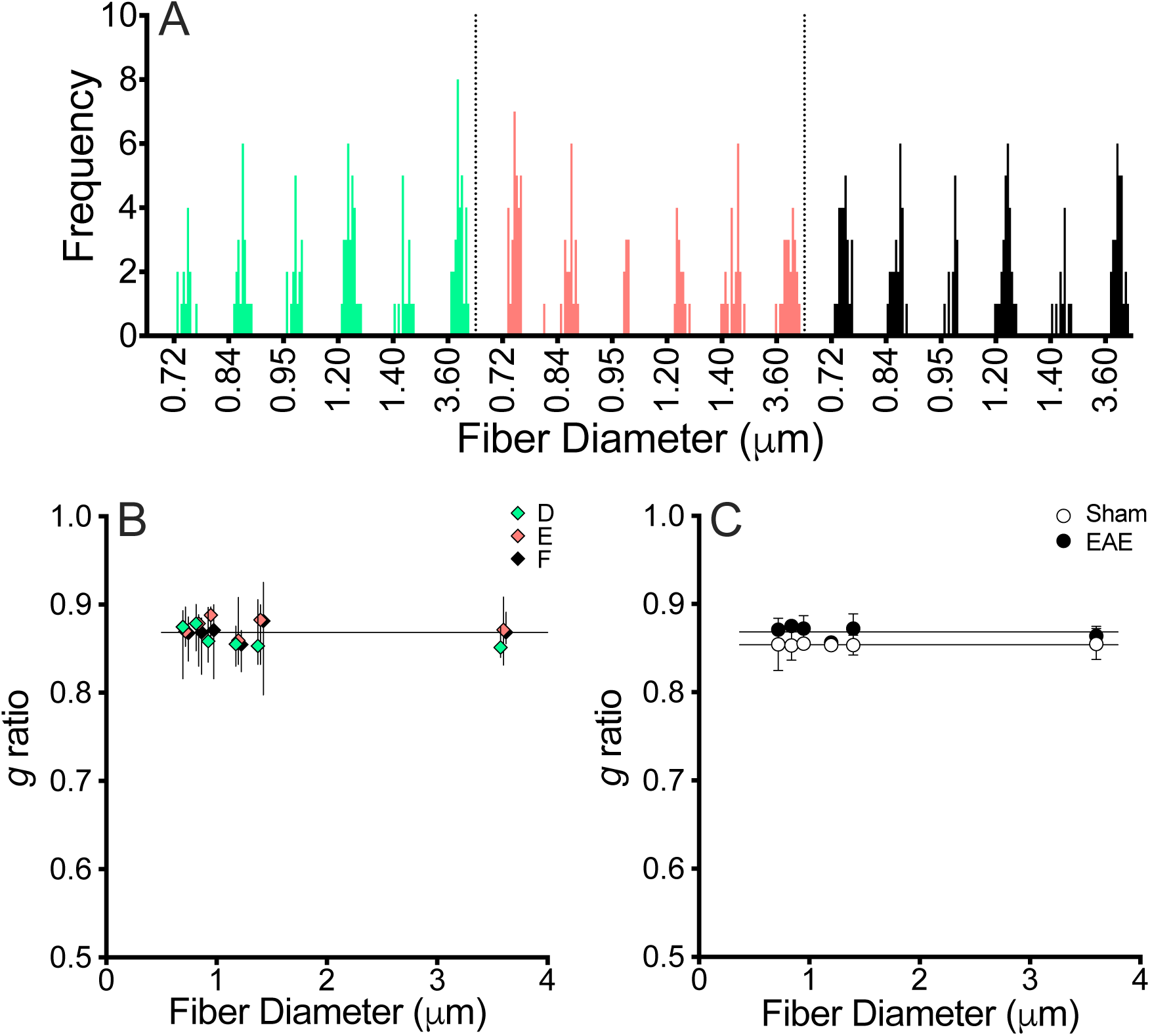
*g* ratio scatterplot quality control and data reduction for EAE treated mice. Scatterplot data from mice D-F (from Fig. 4C) sorted by fiber diameter (ascending order) and binned into six clusters with similar numbers of *g* ratios per cluster (the same cluster boundaries are used for each mouse). **A.** Histograms show *g* ratios in most clusters are unimodal with normal or skewed distributions and can be summarized as medians ±95% CI for each mouse. **B.** The summary data plotted as a relation of the maximum fiber diameter boundary in each cluster for comparison with the horizontal line (grand average for the cohort, Fig. 4C). The data for mice D and F are shifted left and right on the *x*-axis, respectively, to show all the 95% CIs. **C.** Corresponding cluster means mice D-F are averaged (mean ±S.D.). Mixed-effects analysis with Geisser-Greenhouse correction indicates there are no significant differences between clusters (F_(1.24,2.48)_ = 1.79, ε = .25, P = .30), indicating *g* ratio is uncorrelated with fiber diameter. This result is consistent with the direct proportionality of the axon-fiber diameter relation (Fig. 2C). Finally, cluster means for mice A-C (Fig. 3C) are included for comparison with the EAE mice. Mixed-effects analysis for between-treatment differences indicates no differences in *g* ratios for the EAE cohort compared to controls (F_(1,4)_ = 3.28, P = .144).

### Familial hypomyelination in adult mice disrupts axomyelin interactions

In view of the advantages in sensitivity and success of our analysis pipeline over other analysis approaches, we investigated optic nerve data from a second preclinical model with non-immune mediated pathobiology (Griffiths et al., 1990), the *rumpshaker* (*rsh*) mouse. Pathology in this genetic mutant phenocopies developmental hypomyelination in humans (Kobayashi et al., 1994), and harbors the same missense mutation in the *PROTEOLIPID PROTEIN 1* gene (*PLP1*). The *g* ratio scatterplots (Fig. 6A and 6C) from previously published (Southwood et al., 2017) seventy day old wild type (WT) and *rsh* mice (see Methods) are overall similar to the EAE study (Figs 2A, 4A) with regression slope and *y*-intercept instability, but the *g* ratio range in the *rsh* data (particularly mouse B) is considerably more variable than in the other mouse cohorts (Fig. S1F versus S1C).

**Figure 6.**
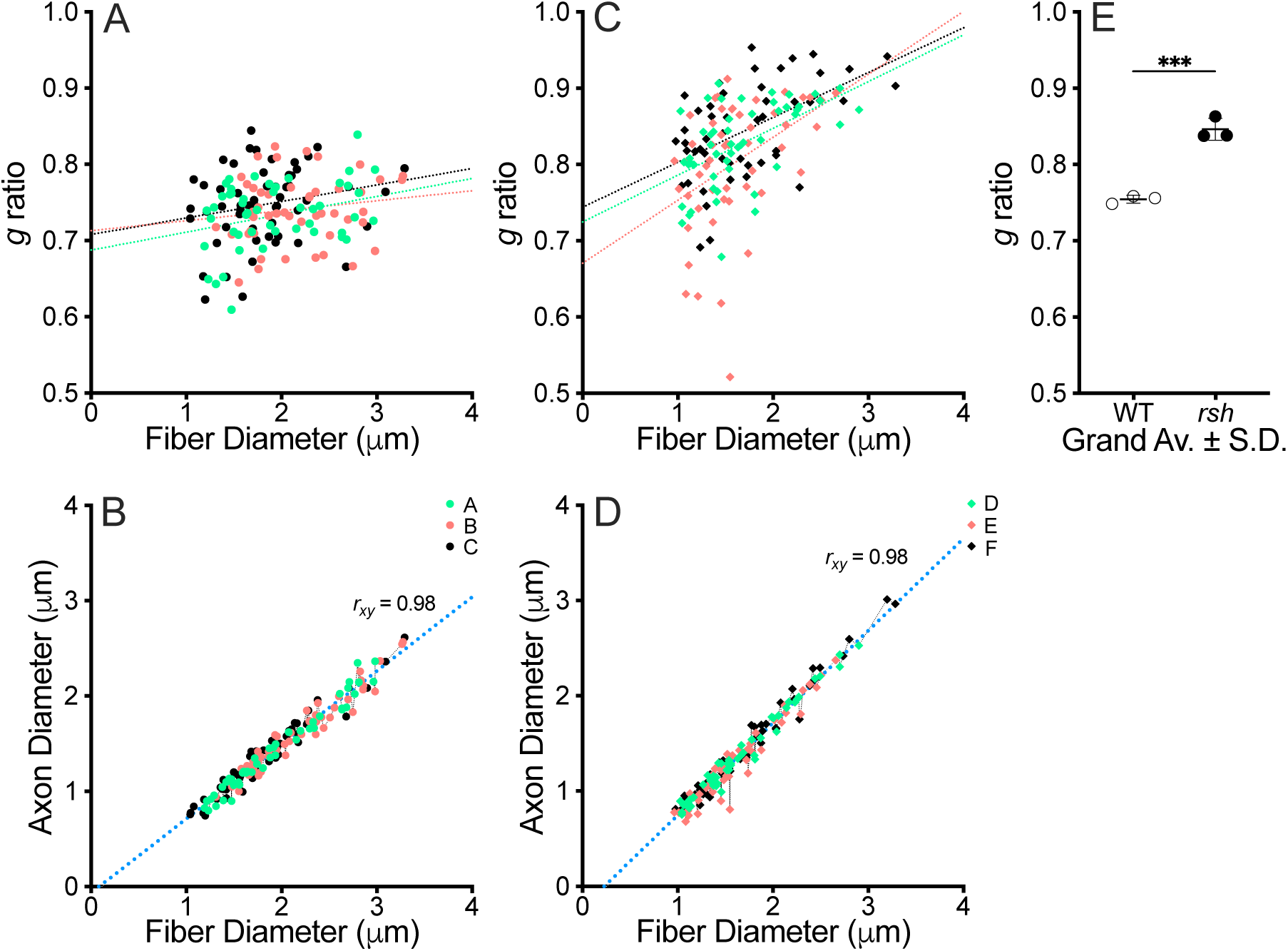
optic nerve *g* ratio versus fiber diameter scatterplots from male WT and *rsh* mice. **A., C.** *g* ratio scatterplots for WT mice and littermate *rsh* mice. Simple linear regression fits are similar in each cohort but differ substantially between the two genotypes. Most of the data are clustered between fiber diameters of 1 – 2.5μm and *g* ratios between .65 – .9. **B., D.** Correlation plots between axon and fiber diameters for the WT and *rsh* cohorts, respectively. The Pearson’s correlation coefficient and Deming regression fit of the control data (B) demonstrates a very strong linear relation (*r_xy_* = .98). The *y*-intercept = −.074 passes very close to the Origin (F_(2,160)_ = 1.91, P = .15) indicating a directly proportional relation. For the *rsh* cohort (D), *r_xy_* = .98 also indicates a strong linear relation between the variables; however, the Deming regression fit shows the *y*-intercept is far from the Origin (−.231 75% CI: [−.308, −.154]). **E.** Grand averages (mean ±S.D.) for each cohort computed from the medians for each mouse in the scatterplots (A) and (C), respectively: WT, .74 ± .01; *rsh*, .85 ± .01. An unpaired t-test shows *g* ratios are increased in the *rsh* cohort (P < .0005, *n* = 3).

Such variability notwithstanding, axon versus fiber diameter plots from the WT and *rsh* cohorts are very strongly correlated and indicative of linear relations (Fig. 6B, 6D), which are confirmed in Figs S4 and S5. Deming linear regression fit to the WT data passes very close to the Origin, suggesting close to direct proportionality (*y*-intercept = −0.065 95% CI: [−0.117, −0.014]). In contrast, the *y*-intercept for the *rsh* data is different from the Origin (*y*-intercept = −0.216, 95% CI: [−0.265, −0.166]), suggesting pathogenic processes in these mutants may impact axomyelin interactions. Indeed, comparison of the grand average *g* ratios in Fig. 6E shows a statistically-significant increase indicative of hypomyelination in the mutants. However, it is unclear if the hypomyelination is across all fiber calibers or is restricted to a subset (e.g. small versus large).

Fiber diameter histogram clusters from the WT and *rsh* cohorts (Fig. 7A and 7B, respectively) show they are unimodal with normal or skewed distributions. There were a relatively small number of measurements per mouse in the original study, so only five fiber clusters are defined. Summary of the clusters (medians ±95% CI) for the WT (Fig. 8A) cohort show they are concordant with the grand average *g* ratio, which is confirmed with a mixed-effects analysis.

**Figure 7.**
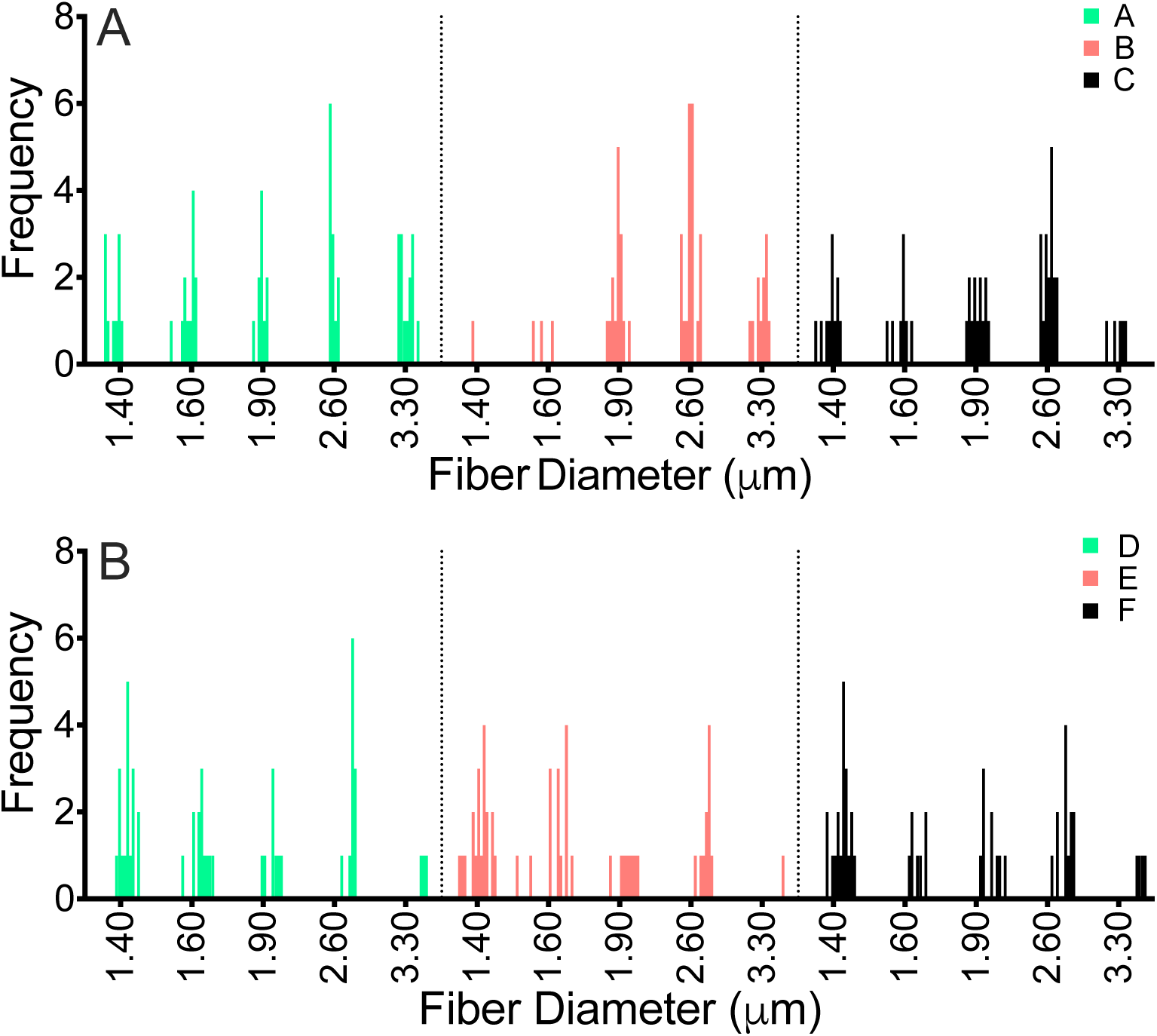
*g* ratio scatterplot quality control for male WT and *rsh* mice. Scatterplot data from mice (from Fig. 6) sorted by fiber diameter (ascending order) and binned into five clusters with similar numbers of *g* ratios per cluster (the same cluster boundaries are used for each mouse). **A.** Histograms from WT mice A-C show *g* ratios in most clusters are unimodal with normal or skewed distributions and can be summarized as medians ±95% CI for each mouse. **B.** Histograms from *rsh* mice D-F show *g* ratios in most clusters are unimodal with normal or skewed distributions and can be summarized as medians ±95% CI for each mouse.

**Figure 8.**
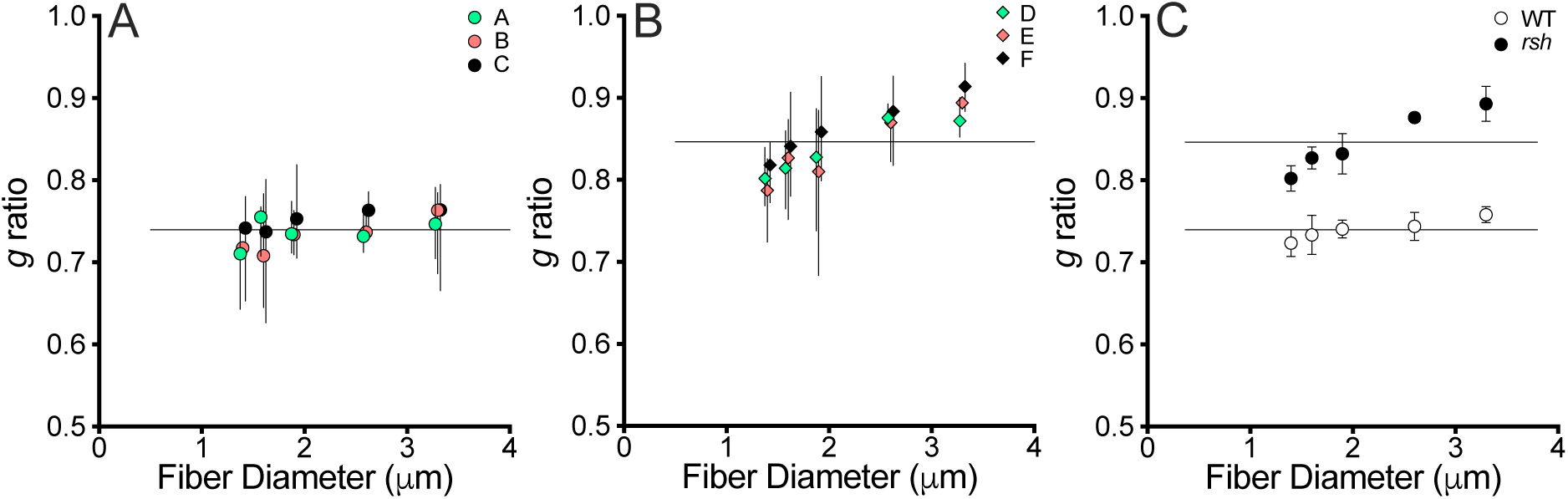
Mixed-effects analysis of WT and *rsh g* ratios from optic nerve. **A.** The summary data from Fig. 7A plotted as a relation of the maximum fiber diameter boundary in each cluster for comparison with the horizontal line (grand average for the cohorts, Fig. 6E). The data for mice A and C are shifted left and right on the *x*-axis, respectively, to show all the 95% CIs. Mixed-effects analysis with Geisser-Greenhouse correction indicates there are no significant differences between clusters (F_(1.30,2.60)_ = 2.55, ε = .33, P = .23). The *g* ratio clusters are somewhat noisy but support a horizontal fit (F_(2,12)_ = 1.91, P = .19), and the confidence interval of the slope is close to zero (.008 < slope < .025). Thus, on balance the *g* ratio is likely uncorrelated with fiber diameter, which conforms to the axomyelin unit model. **B.** The summary data from Fig. 7B plotted as a relation of the maximum fiber diameter boundary in each cluster for comparison with the horizontal line (grand average for the cohorts, Fig. 6E). The data for mice D and F are shifted left and right on the *x*-axis, respectively, to show all the 95% CIs. Mixed-effects analysis with Geisser-Greenhouse correction indicates there are significant differences between cluster means (F_(1.82,3.64)_ = 36.3, ε = .46, P = .004). This result is consistent with the direct proportionality of the axon-fiber diameter relation (Fig. 2C).On the other hand, Tukey’s multiple comparisons test reveals only small differences between a few cluster means (P > .014). Overall, there is weak evidence for a non-horizontal regression fit; thus, *g* ratio is likely correlated with fiber diameter. **C.** Comparison of cluster means for mice A-C and D-F (Fig. 8A and 8B, respectively). Mixed-effects analysis for between-cohort differences indicates strong differences in *g* ratios for the *rsh* cohort compared to controls (F_(1,4)_ = 105, ε = .58, P = .0005).

The same analysis for the *rsh* cohort (Fig. 8B) shows that the grand average *g* ratio line overlaps with most of the 95% CIs for the clusters, but the smallest and largest fibers do not. This suggests a bone fide positive correlation between *g* ratio and fiber diameter, which may be important for understanding the pathobiology. Mixed-effects analysis indicates a statistically significant difference between the fiber diameter clusters (P = 0.004), but post hoc testing reveals only modest differences between some clusters. Thus, the results suggest a weak positive correlation between *g* ratio and fiber diameter such that large diameter fibers may be hypomyelinated relative to the smallest fibers.

Combining Fig. 8A and 8B (Fig. 8C) for mixed-effects analysis yields two important results. First, the principal result is that *g* ratios from *rsh* mice are higher than the controls. Second, error testing between fiber diameter clusters and treatment groups is significant (i.e. testing for interaction between the independent variables). This indicates the relationship between *g* ratio and fiber diameter is influenced by the genotype, which we know from Fig. 8A does not arise from the WT mice and must be associated with the *rsh* cohort. Within-groups difference testing (i.e. comparing fiber diameter clusters within each treatment group) confirms this second result (Fig. 8B). Together, these data enable us to conclude that: (1) axons are hypomyelinated in the *rsh* mice which accords with our conclusion in the original study (Southwood et al., 2017), and (2) the *rsh* pathology may perturb regulatory interactions in the axomyelin unit, where large diameter fibers are comparatively more severely affected than small fibers. This conclusion is unclear in the original study. Thus, the current analysis pipeline provides deeper insight into the pathobiology of the *rsh* mice than we gained previously.

### Equating changes in *g* ratios with changes in myelin thickness

The final tool incorporated into our analysis pipeline is a summary plot that integrates *g* ratio data with a striped estimator for myelin loss or excess (Fig. 9). The processed EAE data from Fig. 5C are shown in panel 9A, with the Sham cohort normalized to 100% myelination according to the grand average *g* ratio. The EAE cohort grand average is slightly above the controls and is not statistically significant (< 10% difference).

**Figure 9.**
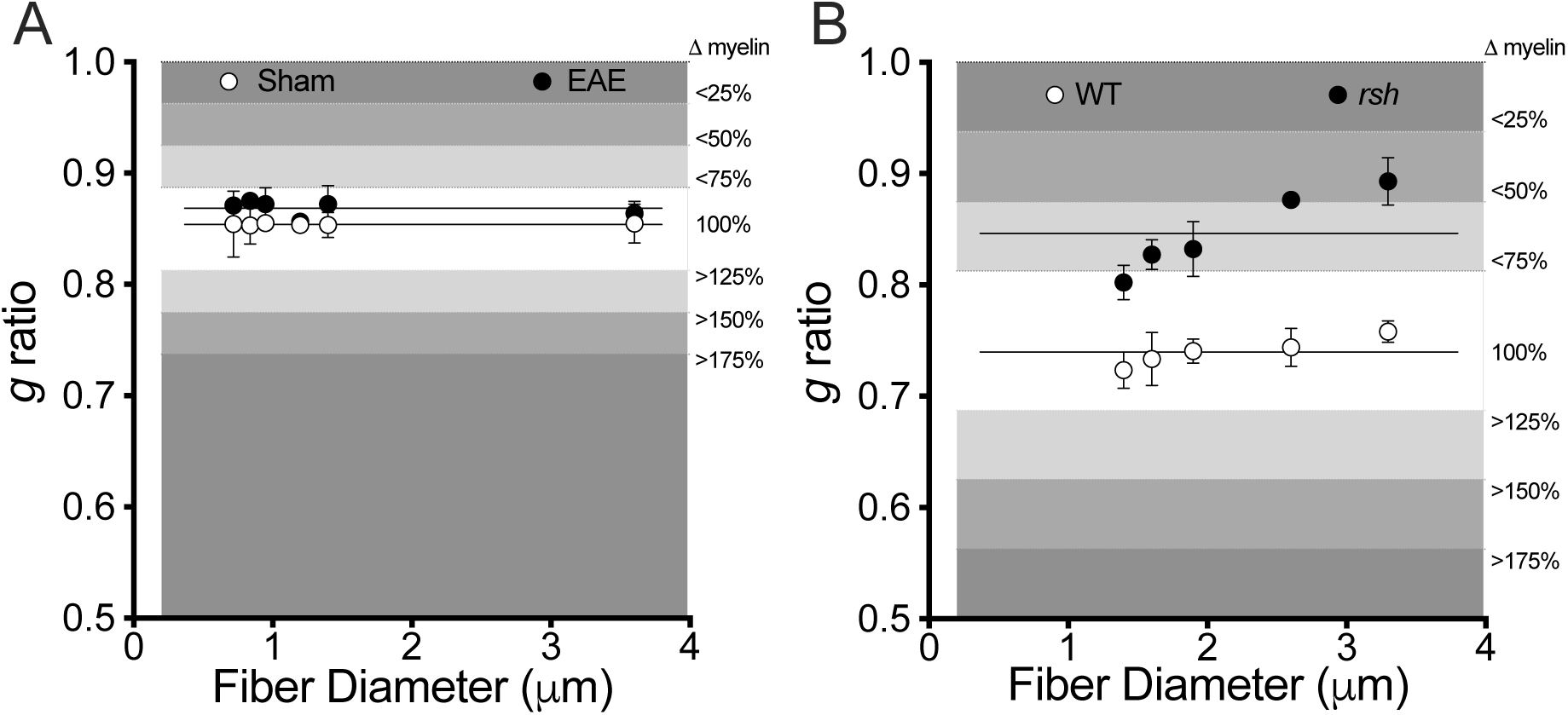
biological impact of in *g* ratio changes in EAE and hypomyelinating disease. The background of the scatterplots is a series of horizontal shaded quartile zones (see key on the right) centered on the grand average for the Sham cohort, which delineate myelin loss (above the grand average) or gain (below). **A.** Scatterplot from Fig. 5C overlaid onto the background. The Sham cluster means show that under physiological conditions myelination across fiber calibers varies minimally. The same is true for the EAE cohort, but there may be a 10-15% decrease in myelin thickness. **B.** Scatterplot from Fig. 8C overlaid onto the background. The grand average *g* ratio in WT littermates is significantly lower than for the Sham mice (Fig. 9A), which likely reflects large differences in the processing of samples for TEM between each study. Hypomyelination in the *rsh* cohort is extensive across all fiber calibers compared with the controls. However, there may be differences in the extent of hypomyelination, with small fibers having about 25% less myelin while myelin loss is greater than 50% in the largest fibers.

In panel 9B, WT data from the *rsh* study (Fig. 8C) also are centered at 100% according to the grand average. The *rsh* data are considerably higher there is a statistically significant 20-55% reduction in myelin thickness. This reduction may be fiber diameter-dependent with the largest fibers being relatively more hypomyelinated. Assuming the positive regression slope through these data is real, the smallest myelinated fibers in these mice would have a predicted *g* ratio = 0.76. This is comparable to the grand average *g* ratio of the WT cohort, suggesting normal myelination. Finally, the grand average *g* ratio for the *rsh* cohort equates to an average 40% myelin loss across all fiber calibers which is comparable to a previously published estimate of 50% loss in cervical spinal cord from 1 month old *rsh* mice (Griffiths et al., 1990).

## DISCUSSION

Structural studies of the axomyelin unit in CNS white matter tracts over recent decades are replete with scatterplots of *g* ratio versus axon diameter analyzed using (least squares) simple linear regression (akin to Figs 2A, 4A, 6A and 6C). In many if not most cases, the slopes and *y*-intercepts are unstable between different animals in the same treatment group. Such variability, and the absence of a predictive model of the axomyelin unit (see accompanying article), can prompt ambiguous or ad hoc interpretations, particularly for control versus disease or treatment comparisons. Nevertheless, the imperative to persist with such experiments despite unclear interpretations is endemic to the field. To stem this vast tide of unrewarded effort, we have revisited the major insights afforded to us over a century of work, to establish a predictive model of the axomyelin unit and a data analysis pipeline that can be used to understand normal physiology and interpret pathobiology in meaningful ways.

In the current study, the axomyelin unit model avoids many of the most common problems associated with interpreting *g* ratio plots. The solution to such problems notwithstanding, a cautionary note is warranted. The analysis pipeline includes a series of steps, which if misused, can bias the analysis outcome. The most crucial of these is in choosing fiber diameter ranges to bin *g* ratios into clusters for histograms. An unbiased choice of fiber diameter ranges relies on being blinded to the outcome of the analysis pipeline, which carries the usual 5% risk (i.e. in accord with the P < 0.05 level of significance) of making type I or II statistical errors. However, iterative cycles of binning fiber diameters until a visually- or statistically-desirable result is achieved is analogous to cherry picking data, or P value hacking, which increases the risk of type I or II errors and misleading interpretations of the analysis.

A common misconception in *g* ratio analyses is that functional changes in the axomyelin unit can be inferred from differences in *g* ratios between controls and diseased, treated or mutant cohorts; in particular, that higher *g* ratios signify lower CVs. However, we know from Rushton’s analysis (1951) there is only a small (5-10%) difference in CV between a fiber (e.g. 1μm diameter) with *g* ratio = 0.75 and another fiber of the same caliber with *g* ratio = 0.85. This seems counterintuitive but it emerges from the fact that, for fibers of the same caliber, slower CV caused by a thinner sheath can be essentially compensated by faster CV from a larger caliber axon. In brief, myelin thickness and axon diameter are counterbalancing.

Interpreting *g* ratio fluctuations by asserting that subtle structural changes cause significant functional changes, arose from early electrophysiological and morphological studies in PNS proposed by Gasser and Grundfest (1939), who proposed a linear relationship between CV and axon caliber. But this relationship also holds for different caliber fibers with the same *g* ratio. In reality, *g* ratio measurements for a given fiber diameter vary appreciably (e.g. Fig. 2A, 0.65 < *g* ratio < 0.85 for most fibers) with minimal functional consequence (Rushton, 1951). And we know from clinical studies that CVs in PNS nerves need to decline several fold (i.e. major changes) before they are clinically relevant (Saporta et al., 2012). Thus, the Gasser and Grundfest CV-axon caliber relationship is too narrow a perspective to account for the observed associations between structure and function.

Establishing the axomyelin unit model and plotting *g* ratios as a relation of fiber diameter draws strength from at least four sources: 1) a framework (reference model under normal physiology) for *g* ratio analyses, based on the direct proportionality between axon and fiber diameters, that constrains data interpretation; 2) the independence of *g* ratio and fiber diameter, which simplifies statistical analysis; 3) maximizing the concordance of data analysis with the underlying assumptions of statistical tests; 4) robust statistics that support inferences and insights from normal physiology into disease mechanisms.

For example, given a constant fiber diameter, it is readily apparent that changes in *g* ratio should have modest or negligible effects on CV because of the myelin thickness-axon diameter counterbalance. Indeed, this may explain most of the *g* ratio ranges reported in countless studies (e.g. 0.7 – 0.8 in Fig. 6A). Thus, if there is little consequence to function, perhaps there is little need for tight regulation of myelin thickness, at least within the confines of normal physiology. Consequently, *g* ratio fluctuations may reflect normal biological variation or compensatory processes under mildly non-physiological conditions, and only large changes in fiber caliber distributions or *g* ratios (e.g. loss of internodes) may be relevant to disease.

Some of the most important uses of *g* ratio versus fiber diameter plots include determining if axomyelin interactions are preserved/disrupted/restored during demyelination/remyelination, pharmacological interventions or normal development. To assist in these and other analyses of white matter function, we have implemented the widely-accepted property of between *g* ratio versus fiber diameter independence, which is valid both experimentally and theoretically for large myelinated axons (Rushton, 1951) and is corroborated herein for small CNS fibers. Together with the statistically-robust axomyelin unit mode, our statistical pipeline provides sensitivity and interpretability for observable changes.

In the current study, we examine two disease cases for changes to myelin *g* ratios. In EAE mice, changes in optic nerve *g* ratios are not detectable. In another published study, subtle functional changes in the CAPs were observed (Zerimech et al., 2023); however, it is difficult to determine how disease severity and course impact the results (e.g. acute versus chronic disease). Disease in EAE mice contrasts with *rsh* in many respects. Disease is *X*-linked (versus acquired), cell autonomous (versus non cell autonomous), begins perinatally (versus adult onset), affects all oligodendrocytes (versus disseminated but likely incomplete) and is chronic (versus acute or transient). In this light, it is not surprising there might be substantial pathophysiological differences between the disease states. Arguably, the most distinct feature of the *g* ratio plot for *rsh* mice is an apparent dependence of *g* ratio on fiber diameter, where large fibers bear the brunt of hypomyelination while the smallest fibers may be close to normally myelinated.

Considering the likelihood that individual oligodendrocytes myelinate both large and small axons, a disproportionate impact on large fibers may reflect subcellular disruption of cargo supply routes from the cell body. For example, ER-Golgi complexes cluster at the base of cell processes, which elaborate myelin sheaths (Southwood et al., 2016). Thus, chronic pathology and locally disrupted cell metabolism (Southwood et al., 2002) may not respond to nonselective myelin enhancing therapeutics and may actually exacerbate pathology. Indeed, such an outcome has been previously documented for *PLP1* mutations (Schneider et al., 1995; Gow and Lazzarini, 1996). Instead, downregulating *Plp1* gene expression to reduce mutant protein synthesis, or lowering general oligodendrocyte metabolism to minimize mutant PLP1 accumulation may be better approaches (Gow and Lazzarini, 1996; Southwood et al., 2002).

Finally, we restate our goals that the purpose of establishing the axomyelin unit model is to derive an explicit framework for hypothesis testing under normal physiologic conditions (the reference state), which can be extended to interpret the impact of disease. Inherent in these goals are specific methodological considerations and limitations. For example, measurement of *g* ratios in controls must be performed with a sufficiently high magnification of 10-15,000x (< 3nm/pixel) to avoid the Rayleigh diffraction limit (Schnepp and Schnepp, 1971; Price and Sprich, 1975) and measurements should be confined to fibers that are largely free of morphological artifacts. The same principles should be upheld for disease states, even if myelin damage is considered bone fide pathobiology (rather than processing artifact). This standard is critical for complying with the tenets of the *g* ratio, defined as the axon diameter : fiber diameter in a region of compact internodal myelin. Thus, myelin swelling, blistering, infolding, redundant sheaths and tissue-processing artifacts have been observed in many diseases tissues (Southwood et al., 2004; Luchicchi et al., 2021; Joost et al., 2022), but are more appropriately quantified using assays other than *g* ratio plots.

## Conclusion

Undoubtedly, the most important aspect of the current study is the formal adoption of an explicit model of the axomyelin unit under physiologic conditions. This model can be used to make predictions about structure and function (many remain to be explored in future studies), thereby building on the foundations from luminaries in the myelin field with more than a century of careful observations and analyses in many species. From this reference model, we have developed an analysis pipeline, with adequate and appropriate statistical methods, and demonstrated the approach to assess and interpret pathobiology in two disease states in mice, one acquired and the other congenital. Important next steps will include the use of this pipeline to explore properties of the myelin internode in multiple species, including humans, and determine its usefulness in mammalian and possibly vertebrate physiology and pathobiology.

## Acknowledgements

This work was supported by grants to A.G. from Wayne State University (Boost Award) and to D.L.F. (BX002625 and 14S-RCS-003 from the Department of Veterans Affairs). We thank the organizers of the 53^rd^ American Society for Neurochemistry annual meeting, Lexington, Kentucky, USA, March 18-22, 2023 for the opportunity to present this work in the colloquium entitled “History, Theory, Applications, Use, and Misuse of the *g*-ratio”. A.G. contributed the *rsh* study data and designed, performed and interpreted the analyses and wrote the manuscript; J.L.D. D.L.F and A.B. contributed the EAE study data and with A.B., were involved in discussions for the manuscript, editing, planning and contributing to the success of the colloquium. The authors declare there are no conflicts of interest.

