## Supplemental Figures 1-5 for "A Statistically-Robust Model Of The Axomyelin Unit Under Normal Physiologic Conditions With Application To Disease States"

Figure S1 – frequency histograms for the EAE and *rsh* studies

**A., B., C.** frequency histograms of the Sham and EAE cohorts (Dupree et al., 2015) for fiber diameter (A) and axon diameter (B) show leftward shifts in the EAE compared to the controls. The *g* ratios (C) are right-shifted, indicating overall substantially thinner myelin in the EAE cohort. The fibers measured in this study were randomly selected and represent unbiased distributions of fibers within the optic nerves of the mice. **D., E., F.** frequency histograms of the WT and *rsh* cohorts (Southwood et al., 2017) for fiber diameter (D) and axon diameter (E) show leftward shifts for the spread of diameters in *rsh* compared to controls, more so in the fibers than the axons. The *g* ratios (F) are right-shifted indicating overall thinner myelin in the *rsh* cohort. Importantly, the fibers measured in this study were selected for size distribution, and do not represent unbiased distributions of fibers within the optic nerves of the mice.

Figure S2 – Statistical and biological constraints on axon-fiber diameter plots demonstrates a direct proportionality relation for the Sham (EAE) cohort

**A.** Summary of a Box-Cox lambda transformation series for axon diameters from mouse A plotted against cognate fiber diameters (Fig. 2C). The transforms are fit using simple linear regression, and extrapolated to the *x*- and *y*-axes to estimate several parameters that identify the most likely axon-fiber diameter relation. **B.** Pearson's correlation coefficients for individual regression fits (mice A-C, Fig. 2C) plotted against the corresponding Box-Cox lambda values. The maximum $r_{xy}$ value, for $\lambda = 1$, is the most likely axon-fiber diameter relation as confirmed by mixed-effects analysis with Geisser-Greenhouse correction ($F_{(1.23,2.65)} = 1010$, $\varepsilon = .18$, $P = .0003$) following Fisher's Z transformations of the $r_{xy}$ values. Post hoc Dunnett's tests show that all other values of lambda yield inferior correlations ($P < .028$). **C.** The *x*- and *y*-axis intercepts ±95% CI for the extrapolated regression fits in (A) plotted against the Box-Cox lambda values. Biological constraints on these intercepts exclude regression fits for all *x*-intercepts $\leq 0$ and *y*-intercepts > 0, i.e. for all transformations of $D_A$ where $\lambda < 1$. **D.** The constraints of myelinated fibers mandate that the regression slope < 1 (i.e. $D_F > D_A$), which excludes regression fits where $\lambda > 1$.

Figure S3 – Several statistical and biological constraints on axon-fiber diameter plots demonstrates a direct proportionality relation for the EAE cohort

**A.** Summary of a Box-Cox lambda transformation series for axon diameters from mouse D in Fig. 4 plotted against cognate fiber diameters. The transforms are fit using simple linear regression, and extrapolated to the *x*- and *y*-axes to estimate several parameters that identify the most likely axon-fiber diameter relation. **B.** Pearson's correlation coefficients for individual regression fits (mice D-F, Fig. 4B) plotted against the corresponding Box-Cox lambda values. The maximum $r_{xy}$ value, for $\lambda$ = 1, is the most likely axon-fiber diameter relation as confirmed by mixed-effects analysis with Geisser-Greenhouse correction ($F_{(1.63,3.73)} = 53.4$, $\varepsilon = .23$, $P = .002$) following Fisher's Z transformations of the $r_{xy}$ values. Post hoc Dunnett's tests show that no other values of lambda yield superior correlations ($P > .05$). **C.** The *x*- and *y*-axis intercepts ±95% CI for the extrapolated regression fits in (A) plotted against the Box-Cox lambda values. Biological constraints on these intercepts exclude regression fits for all *x*-intercepts $\leq 0$ and *y*-intercepts $> 0$, i.e. for all transformations of $D_A$ where $\lambda < 1$. **D.** The constraints of myelinated fibers mandate that the regression slope < 1 (i.e. $D_F > D_A$), which excludes regression fits where $\lambda > 1$.

Figure S4 – Several statistical and biological constraints on axon-fiber diameter plots demonstrates a direct proportionality relation for the WT (*rsh*) cohort

**A.** Summary of a Box-Cox lambda transformation series for axon diameters from mouse A in Fig. 6 plotted against cognate fiber diameters. The transforms are fit using simple linear regression, and extrapolated to the *x*- and *y*-axes to estimate several parameters that identify the most likely axon-fiber diameter relation. **B.** Pearson's correlation coefficients for individual regression fits (mice A-C, Fig. 6B) plotted against the corresponding Box-Cox lambda values. The maximum $r_{xy}$ value, for $\lambda = 1$, is the most likely axon-fiber diameter relation as confirmed by mixed-effects analysis with Geisser-Greenhouse correction ($F_{(1.39,2.77)} = 54.9$, $\varepsilon = .20$, P = .006) following Fisher's Z transformations of the $r_{xy}$ values. Post hoc Dunnett's tests show that no other values of lambda yield superior correlations (P > .05). **C.** The *x*- and *y*-axis intercepts ±95% CI for the extrapolated regression fits in (A) plotted against the Box-Cox lambda values. Biological constraints on these intercepts exclude regression fits for all *x*-intercepts ≤ 0 and *y*-intercepts > 0, i.e. for all transformations of $D_A$ where $\lambda < 1$. **D.** The constraints of myelinated fibers mandate that the regression slope < 1 (i.e. $D_F > D_A$), which excludes regression fits where $\lambda > 1$.

Figure S5 – Several statistical and biological constraints on axon-fiber diameter plots demonstrates a direct proportionality relation for the *rsh* cohort

**A.** Summary of a Box-Cox lambda transformation series for axon diameters from mouse D in Fig. 6 plotted against cognate fiber diameters. The transforms are fit using simple linear regression, and extrapolated to the *x*- and *y*-axes to estimate several parameters that identify the most likely axon-fiber diameter relation. **B.** Pearson's correlation coefficients for individual regression fits (mice D-F, Fig. 6D) plotted against the corresponding Box-Cox lambda values. The maximum $r_{xy}$ value, for $\lambda = 1$, is the most likely axon-fiber diameter relation as confirmed by mixed-effects analysis with Geisser-Greenhouse correction ($F_{(1.20,2.40)} = 47.7$, $\varepsilon = .17$, $P = .012$) following Fisher's Z transformations of the $r_{xy}$ values. Post hoc Dunnett's tests show that no other values of lambda yield superior correlations ($P > .08$). **C.** The *x*- and *y*-axis intercepts ±95% CI for the extrapolated regression fits in (A) plotted against the Box-Cox lambda values. Biological constraints on these intercepts exclude regression fits for all *x*-intercepts $\leq 0$ and *y*-intercepts $> 0$, i.e. for all transformations of $D_A$ where $\lambda < 1$. **D.** The constraints of myelinated fibers mandate that the regression slope < 1 (i.e. $D_F > D_A$), which excludes regression fits where $\lambda > 1$.

Figure S1 –

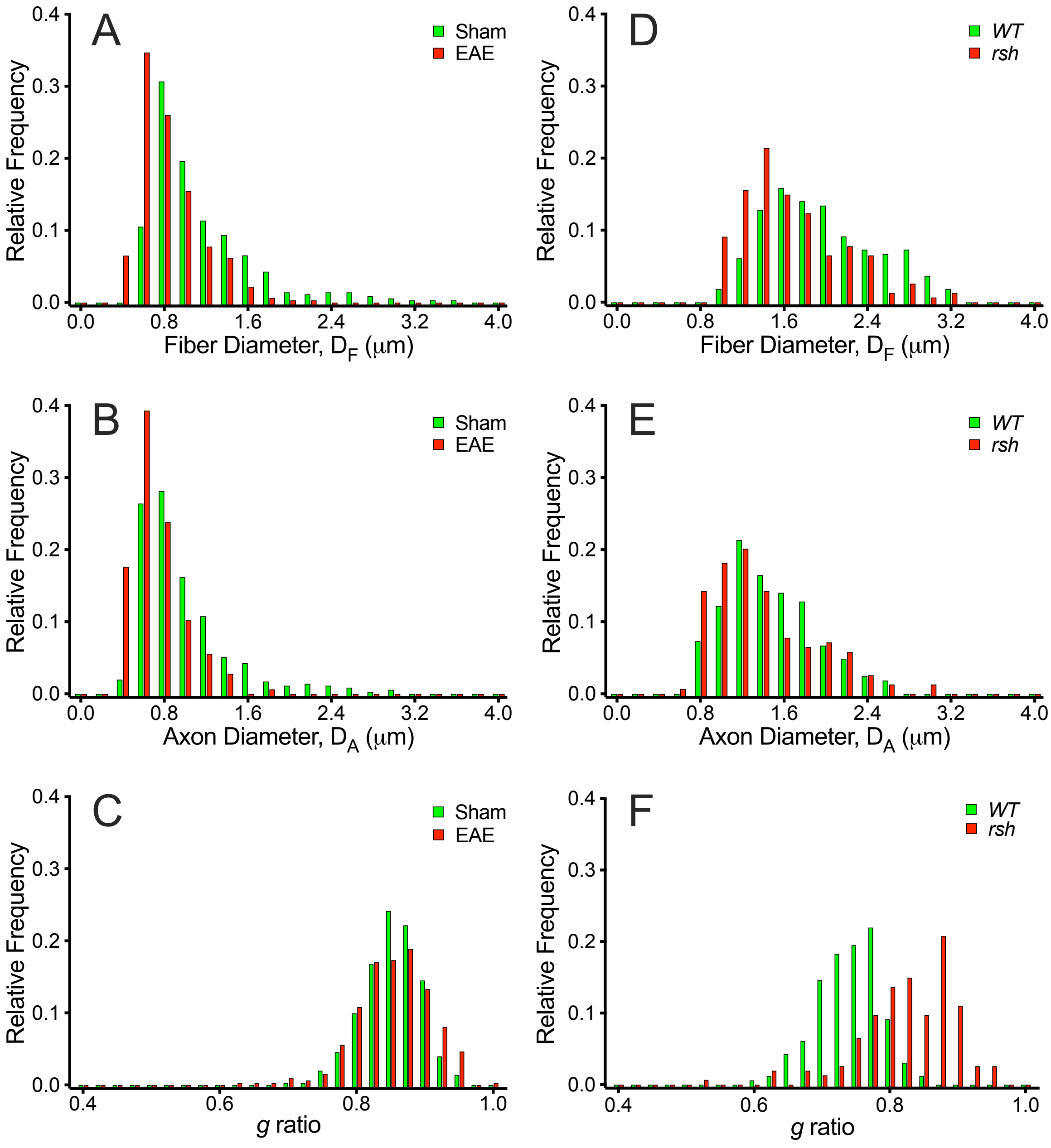
A
Sham
EAE
Relative Frequency
Fiber Diameter, $D_F$ (μm)
B
Axon Diameter, $D_A$ (μm)
C
g ratio
D
WT
rsh
E
F
0.0
0.1
0.2
0.3
0.4
0.8
1.6
2.4
3.2
4.0
0.6
1.0

Figure S2 –

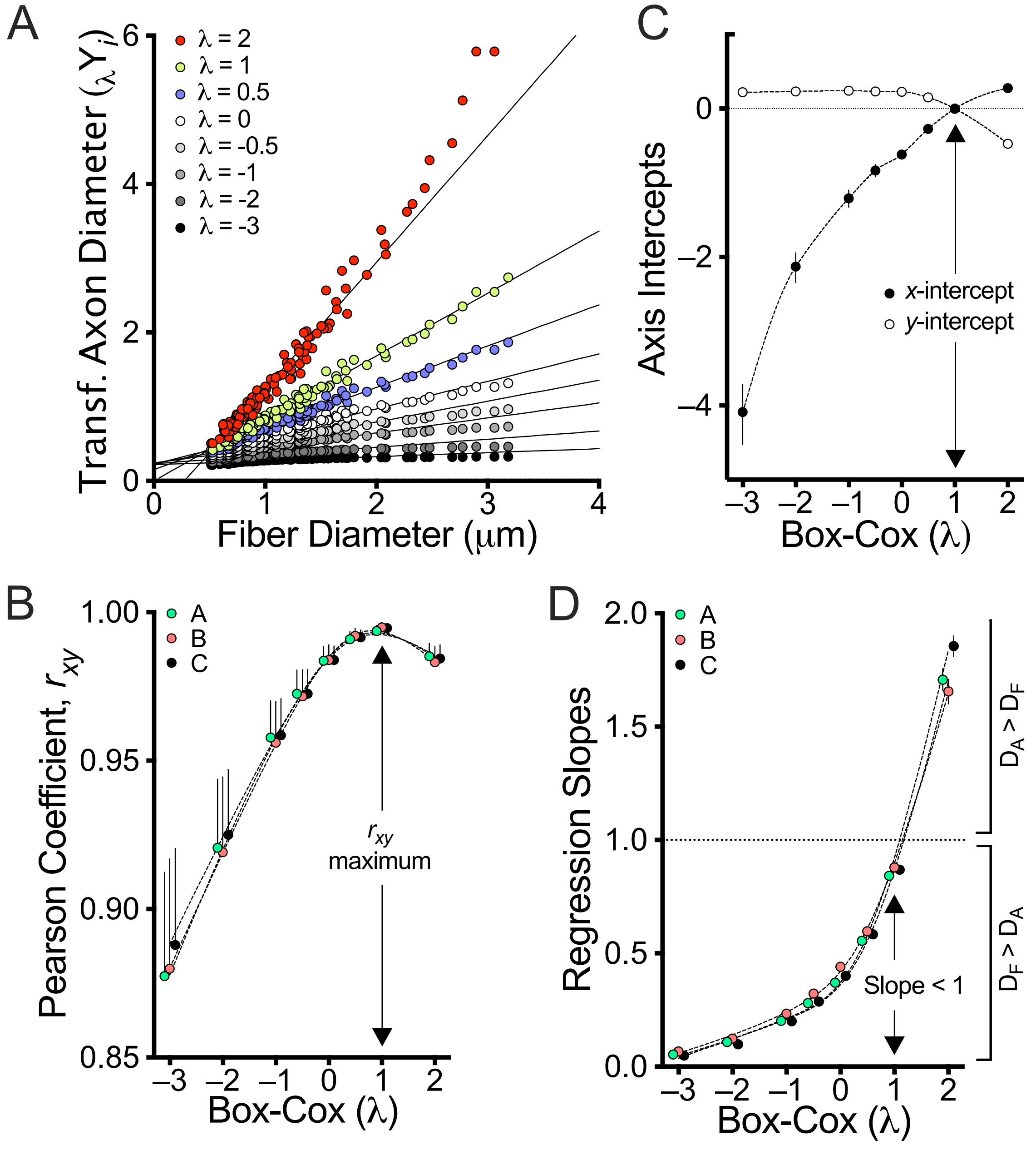

A
Transf. Axon Diameter ($_\lambda Y_i$)
λ = 2
λ = 1
λ = 0.5
λ = 0
λ = -0.5
λ = -1
λ = -2
λ = -3
0
2
4
6
0
1
2
3
4
Fiber Diameter (μm)
C
Axis Intercepts
0
−2
−4
x-intercept
y-intercept
−3
−2
−1
0
1
2
Box-Cox (λ)
B
Pearson Coefficient, $r_{xy}$
1.00
0.95
0.90
0.85
A
B
C
$r_{xy}$
maximum
−3
−2
−1
0
1
2
Box-Cox (λ)
D
Regression Slopes
2.0
1.5
1.0
0.5
0.0
A
B
C
$D_A > D_F$
$D_F > D_A$
Slope < 1
−3
−2
−1
0
1
2
Box-Cox (λ)

Figure S3 –

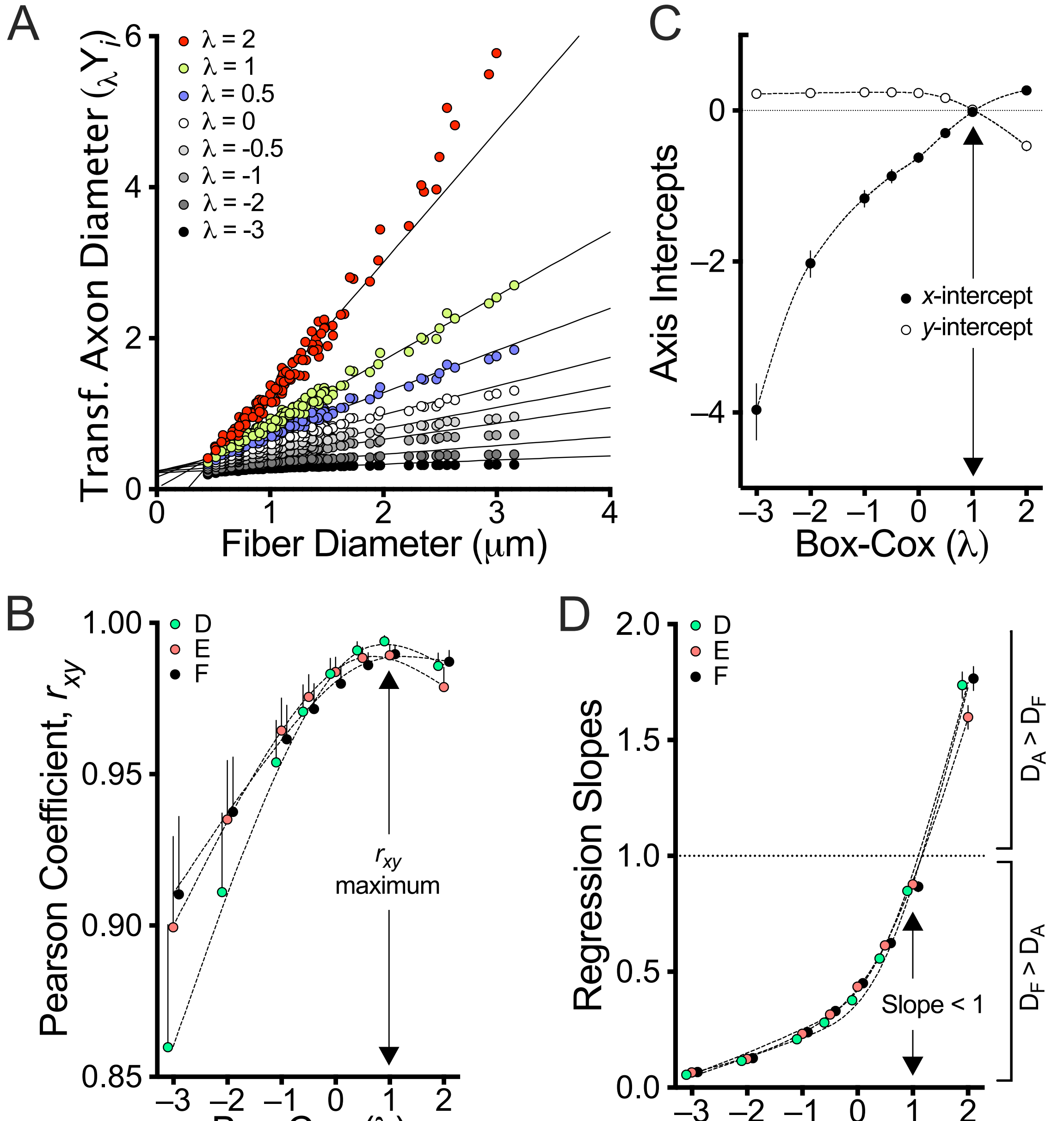

A
Transf. Axon Diameter ($_\lambda Y_i$)
λ = 2
λ = 1
λ = 0.5
λ = 0
λ = -0.5
λ = -1
λ = -2
λ = -3
Fiber Diameter (μm)
C
Axis Intercepts
x-intercept
y-intercept
Box-Cox (λ)
B
Pearson Coefficient, $r_{xy}$
D
E
F
$r_{xy}$
maximum
Box-Cox (λ)
D
Regression Slopes
D
E
F
$D_A > D_F$
$D_F > D_A$
Slope < 1
Box-Cox (λ)

Figure S4 –

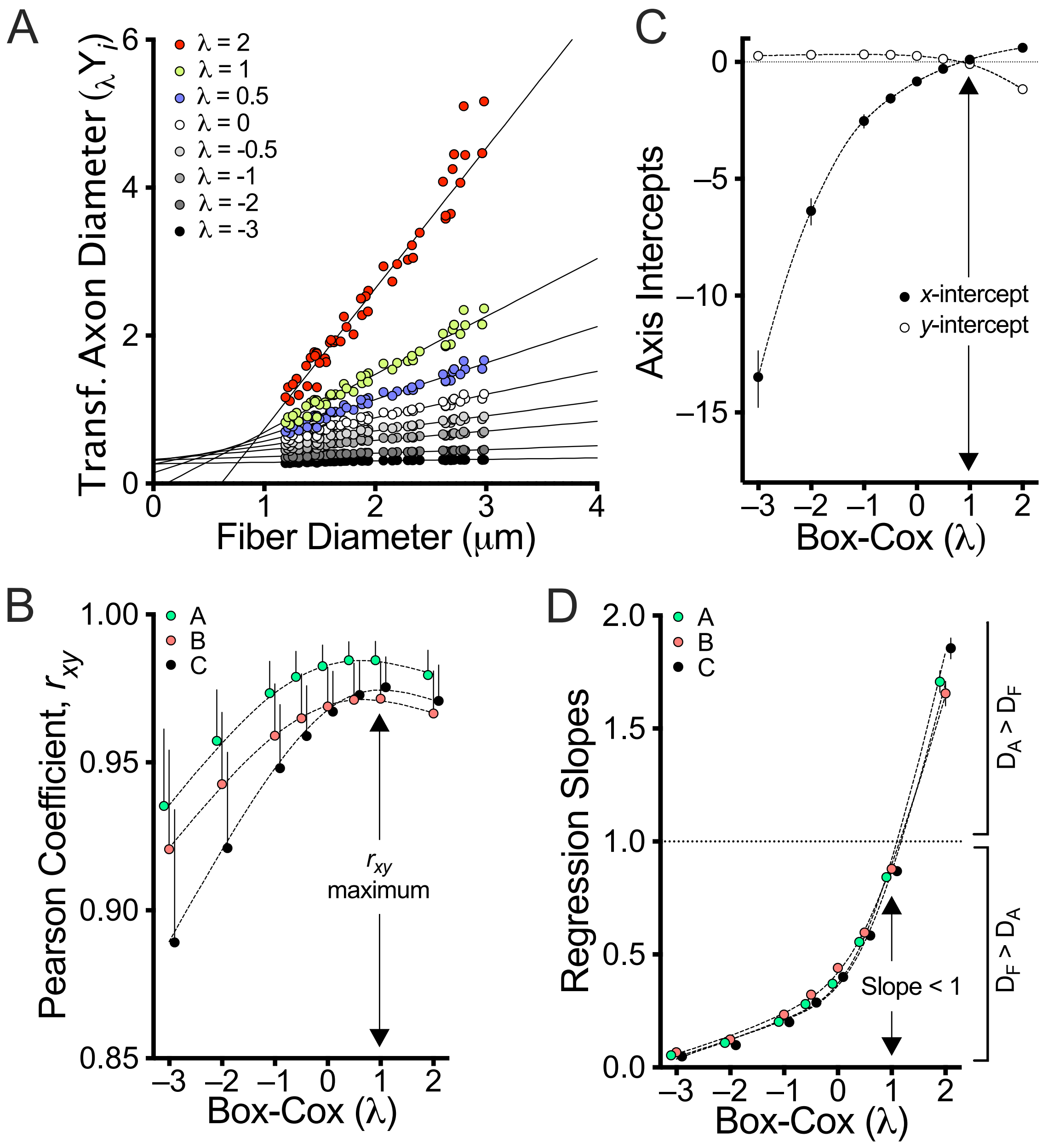
A
Transf. Axon Diameter ($_\lambda Y_i$)
λ = 2
λ = 1
λ = 0.5
λ = 0
λ = -0.5
λ = -1
λ = -2
λ = -3
Fiber Diameter (μm)
C
Axis Intercepts
x-intercept
y-intercept
Box-Cox (λ)
B
Pearson Coefficient, $r_{xy}$
A
B
C
$r_{xy}$ maximum
Box-Cox (λ)
D
Regression Slopes
A
B
C
$D_A > D_F$
$D_F > D_A$
Slope < 1
Box-Cox (λ)

Figure S5 –

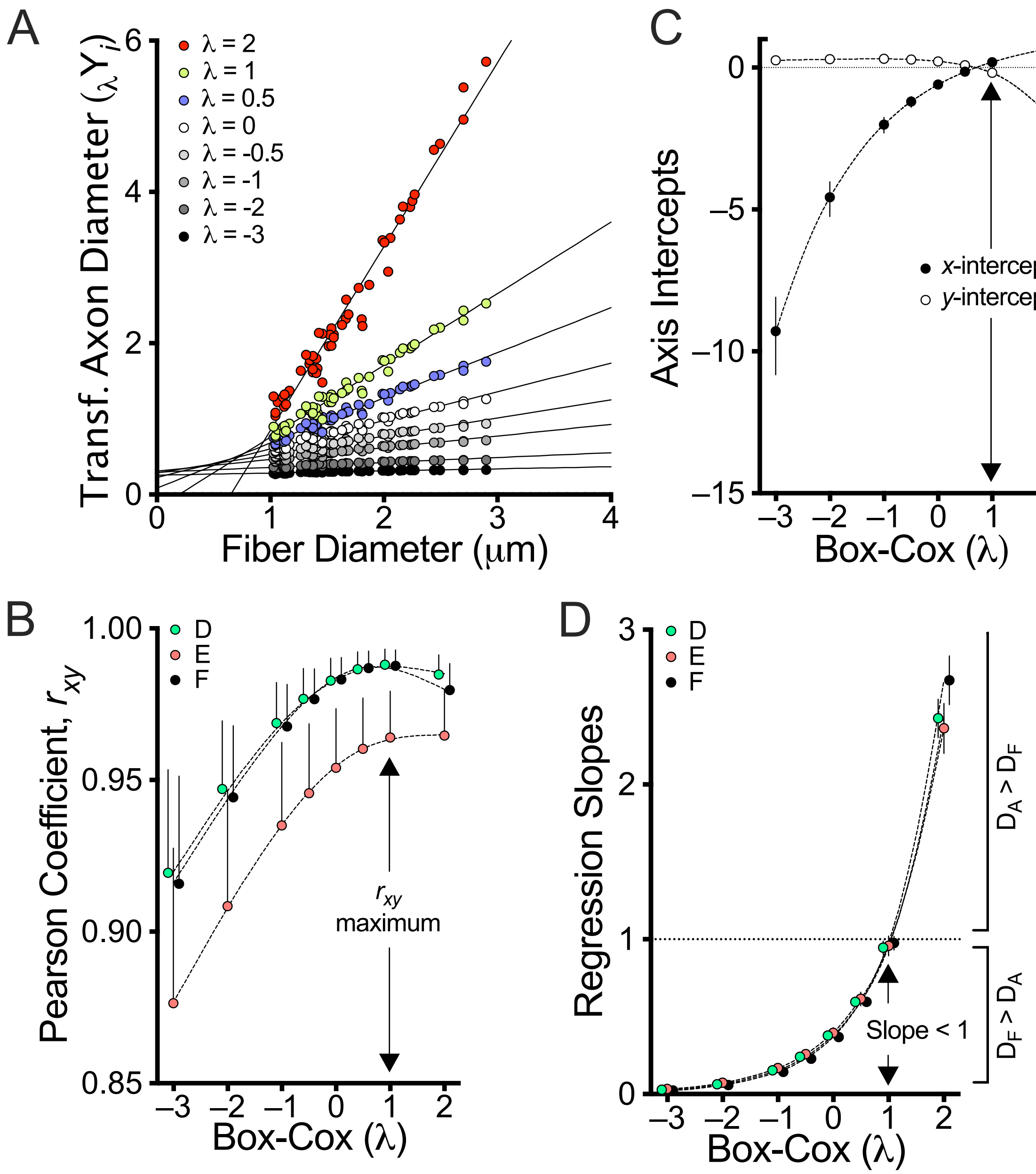

A
Transf. Axon Diameter ($_\lambda Y_i$)
Fiber Diameter (μm)
λ = 2
λ = 1
λ = 0.5
λ = 0
λ = -0.5
λ = -1
λ = -2
λ = -3
B
Pearson Coefficient, $r_{xy}$
Box-Cox (λ)
D
E
F
$r_{xy}$ maximum
C
Axis Intercepts
Box-Cox (λ)
x-intercept
y-intercept
D
Regression Slopes
Box-Cox (λ)
Slope < 1
$D_A > D_F$
$D_F > D_A$
